# Translational T-box riboswitches bind tRNA by modulating conformational flexibility

**DOI:** 10.1101/2024.02.02.578613

**Authors:** Eduardo Campos-Chavez, Sneha Paul, Zunwu Zhou, Dulce Alonso, Anjali R. Verma, Jingyi Fei, Alfonso Mondragón

**Affiliations:** Department of Molecular Biosciences, Northwestern University, Evanston, IL 60208, USA; Institute for Biophysical Dynamics and Department of Biochemistry and Molecular Biology, The University of Chicago, Chicago, IL 60637, USA; Department of Chemistry, Columbia University, New York, NY 10027, USA; Current address: Institute of Molecular Sciences of Orsay, Paris-Saclay University, 91405 Orsay, France (SP); Current address: Biophysics Program and Institute for Physical Sciences and Technology, University of Maryland, College Park, MD 20742 (AV)

## Abstract

T-box riboswitches, paradigmatic non-coding RNA elements involved in genetic regulation in most Gram-positive bacteria, are adept at monitoring amino acid metabolism through direct interactions with specific tRNAs. T-box riboswitches assess tRNA aminoacylation status, subsequently regulating the transcription or translation of downstream genes involved in amino acid metabolism. Here we present single-molecule FRET studies of the *Mycobacterium tuberculosis IleS* T-box riboswitch, a model of T-box translational regulation. The data supports a two-step binding model where the tRNA anticodon is recognized first, followed by interactions with the NCCA sequence. Specifically, after anticodon recognition, tRNA in the partially bound state can transiently dock into the discriminator domain, resembling the fully bound state, even in the absence of the tRNA NCCA-discriminator interactions. Establishment of the NCCA-discriminator interactions significantly stabilizes the fully bound state. Collectively, the data suggests higher conformational flexibility in translation-regulating T-box riboswitches, compared to transcription-regulating ones, and supports a conformational selection model for NCCA recognition. Furthermore, it was found that the conserved RAG sequence is pivotal in maintaining specific interactions with the tRNA NCCA sequence by preventing sampling of an aberrant conformational state, while Stem IIA/B-linker interactions impact the conformational dynamics and the stability of both the partially bound and fully bound states. The present study provides a critical kinetic basis for how specific sequences and structural elements in T-box riboswitches enable the binding efficiency and specificity required to achieve gene regulation.

## Introduction

Protein synthesis, or translation, is an essential cellular activity in all known cellular life. Translation requires aminoacylated tRNAs (charged tRNAs) as substrates, which are generated by aminoacyl-tRNA synthetases. To ensure an adequate supply of charged tRNAs, several regulatory mechanisms have evolved (1). In Gram-positive bacteria, one such mechanism is represented by T-box riboswitches. Riboswitches are *cis*-acting RNA regulatory elements that undergo major structural changes in response to a regulatory signal (2,3). While numerous riboswitches regulate gene expression in response to various small molecule metabolites (3), T-box riboswitches can distinguish and respond to the aminoacylation state of tRNAs (4). An increased ratio of uncharged to charged tRNA indicates a shortage of a specific amino acid or aminoacyl-tRNA synthetase (5). By detecting intracellular concentrations of uncharged tRNAs, T-box riboswitches, located at the 5’ leader region of the mRNAs under their control, can alter the transcription or translation of aminoacyl tRNA synthetase genes or other amino acid metabolism-related genes (6-8). Aside from the ribosome, T-box riboswitches are the only other known RNA capable of both tRNA identity decoding and aminoacylation sensing (9), and thus provide a unique paradigm for gene regulation by two interacting, structured non-coding RNAs (4,6,10,11).

As with other riboswitches, T-box riboswitches are modular and composed of two parts, a ligand binding domain and an expression platform. In the case of T-box riboswitches, tRNA decoding and aminoacylation status discrimination reside in two separate structural domains (**Fig. 1a**). T-box riboswitches have an obligatory Stem I domain that decodes tRNA identity, (12-14); this process is often facilitated by Stem II and Stem IIA/B domains (6,9). In general, Stem I, Stem II, and Stem II A/B comprise the decoding module (**Fig. 1a**), except for glycine T-box riboswitches, where Stem II and Stem IIA/B are absent. To achieve specificity, the tRNA anticodon pairs with a single codon sequence in the T-box RNA, known as the specifier sequence, which is situated in the specifier loop of Stem I. The discriminator domain examines the aminoacylation status of the tRNA and switches the expression output appropriately (9,15) (**Fig. 1b**). The discriminator domain includes Stem III, an antiterminator or antisequestrator region, and the T-box sequence (AGGGUGGNACCGCG), which comprises the 14 most highly conserved nucleotides of the complete T-box RNA (**Fig. 1a**) and is involved in recognition of the NCCA sequence at the 3’ end of the tRNA.

**Figure 1.**
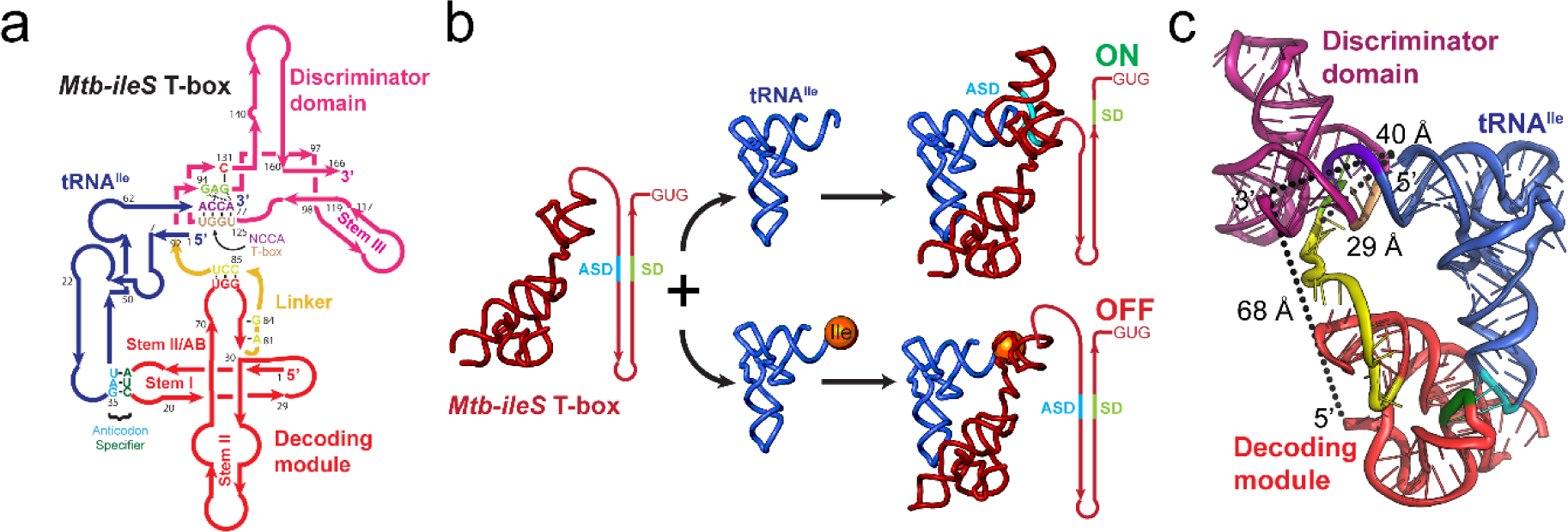
Structure and mechanism of the *Mtb-ileS* T-box riboswitch. **a)** Secondary structure diagram showing the *Mtb-ileS* T-box riboswitch (pink/red) and tRNA^Ile^ (blue). The T-box has two distinct structural domains, the decoding module (red) is involved in recognizing specific tRNAs through anticodon-specifier interactions (cyan, green), whereas the discriminator domain (pink) contains the T-box sequence (brown) involved in recognizing the 3’ NCCA region of the tRNA (purple) and assessing the aminoacylation status. The two domains are joined by a linker region (yellow). The secondary structure diagram is based on the structure of the complex (9). **b)** Proposed mechanism for translation-regulating T-box riboswitches. The T-box decodes the tRNA identity of the tRNA and aminoacylation status at the 3’ end. Depending on the charge state of the tRNA, it exposes or sequesters the Ribosome Binding Site or Shine Dalgarno sequence (SD, green) and the Anti Shine Dalgarno (ASD, cyan) sequence. If the tRNA is uncharged, the 3’ NCCA of the tRNA forms base pairs with the T-box sequence leading to a conformation where the SD region is accessible, and translation is allowed (top). If the tRNA is charged, the interactions with the T-box sequence are precluded, leading to a conformation where the ASD and SD regions interact, the SD is inaccessible, and translation is not allowed (bottom). **c)** Structure of the *Mtb-ileS* T-box riboswitch/tRNA^Ile^ complex (9). The molecules are colored as in **a)**. Approximate distances between the 3’ and 5’ ends of the T-box (69 Å), the 3’ end of T-box and the 5’ end of the tRNA (40 Å), and the 3’ end of the linker region (Δ-Discriminator Mutant) and the 5’ end of the tRNA (29 Å) are shown.

The majority of T-box riboswitches regulate at the transcription level (16-18). In the absence of tRNA or in the presence of charged tRNA, transcription-regulating T-box riboswitches typically adopt an intrinsic terminator structure that promotes premature termination of transcription. Alternatively, binding of an uncharged cognate tRNA promotes the formation of an antiterminator structure to enable transcription readthrough. The *Bacillus subtilis glyQS* T-box riboswitch, as a representative transcription-regulating T-box riboswitch, has been the subject of extensive structural and kinetic studies (12,14,15,19). Single-molecule characterization of the *glyQS* T-box riboswitch established a two-step binding model in which the anticodon of tRNA^Gly^ is recognized first, leading to a partially bound state, followed by binding of the tRNA^Gly^ 3’ NCCA end, leading to a fully bound state (19,20). In addition, an intramolecular conformational change between the decoding and the discriminator domains occurs during the second binding step. In this model, binding of the tRNA^Gly^ ligand by the *glyQS* T-box riboswitch notably follows the directionality of transcription, with the kinetic parameters supporting a co-transcriptional binding and switching model. Specifically, the short lifetime (∼4 sec) of the partially bound state allows rapid sampling of the anticodon sequence and ensures that the tRNA remains bound until synthesis of the discriminator domain occurs. On the other hand, the fast transition into the fully bound state when the uncharged NCCA end is present effectively locks the T-box riboswitch into the antiterminator conformation. Finally, the fully bound state is very stable to allow for transcription readthrough beyond the T-box regulatory sequence.

Translation-regulating T-box riboswitches achieve gene regulation via tRNA-induced conformational changes from a sequestrator conformation, which occludes the Ribosome Binding Site (RBS) sequence, to an antisequestrator conformation, which makes the RBS accessible to the ribosome (**Fig. 1b**) (21). While the core structural domains for anticodon recognition and aminoacylation discrimination are highly similar between the translation and transcription-regulating T-box riboswitches, translation-regulating T-box riboswitches contain distinct structural elements (9). Compared to their transcription-regulating counterparts, translation-regulating T-box riboswitches have a shorter Stem I. Stem I in the *glyQS* T-box riboswitch provides an additional contact with the elbow region of the tRNA that greatly facilitates the first binding step of the tRNA to the riboswitch (14,15). The absence of this critical interaction in translation-regulating T-box riboswitches is partially compensated for by additional interactions provided by Stems II and IIA/B at the anticodon binding pocket (6,9). However, it is unclear whether these structural differences lead to different binding kinetics in translation-regulating T-box riboswitches. In addition, it may be possible for translation-regulating T-box riboswitches to perform several rounds of regulation throughout an mRNA lifetime, rather than being limited to co-transcriptional regulation on the nascent transcript, as is the case for transcriptional T-box riboswitches. Thus, the two types of riboswitches may have evolved different structural dynamics to accommodate their functional differences (22). However, further kinetic studies of the tRNA binding mechanisms and associated conformational dynamics in translational T-box riboswitches are still necessary to elucidate these differences. While recent kinetic characterizations of translation-regulating T-box riboswitches lacking the discriminator domain provide novel mechanistic insights on the conformational landscape of the decoding module and its response to tRNA binding (23,24), it remains unknown how conformational dynamics at the decoding module communicate with the rest of the T-box to coordinate their actions −information which is essential to understand the function of these riboswitches.

Here, the interactions between tRNA and a translational T-box riboswitch are investigated using a combination of single-molecule Förster resonance energy transfer (smFRET), ensemble binding assays, and mutagenesis. The use of a translational T-box riboswitch construct including both decoding and discriminator modules enabled kinetic characterization of the full binding process. The results reveal a two-step tRNA binding process with overall similarities to the binding process observed for transcription-regulating T-box riboswitches, but with distinct kinetic features. Particularly, the data here supports a conformational selection model for binding of the tRNA NCCA end to the translation-regulating T-box. In addition, our data highlights critical structural elements at the decoding and discriminator modules that kinetically facilitate tRNA binding efficiency and specificity. The present work provides a comprehensive picture of the tRNA binding kinetics by a translational T-box riboswitch, thereby illuminating the important functional differences between translation- and transcription-regulating T-box riboswitches.

## Results

### Preparation of fluorophore-labeled *Mtb ileS* T-box riboswitch

The *Mycobacterium tuberculosis ileS* T-box riboswitch (*Mtb ileS* T-box) was chosen as a model system to study translational T-box riboswitches due to the availability of a crystallographic structure of the complex, which encompasses a significant portion of the T-box riboswitch sequence (9). The structural information served as a valuable reference to guide the experimental design. Initially, a construct was designed including a 15-nucleotide extension at the 5’ end of the *in vitro* transcribed T-box RNA, which allowed annealing to a modified DNA probe for immobilization. Purification of this transcript by urea-polyacrylamide gel electrophoresis (urea-PAGE), followed by direct Cy3 labeling at the 3’ end, and *in vitro* refolding in the presence of a biotinylated oligonucleotide resulted in no measurable tRNA^Ile^ binding. This was likely due to the presence of the 5’ extension, which was impeding proper folding of the transcript through *in vitro* refolding. To address this issue, a T-box construct was designed employing an alternative labeling method as follows. The 15-nucleotide extension used for immobilization was appended to the 3’ end of the transcript, downstream of the antisequestrator domain. This construct encompasses Stem I, Stem II, Stem II-A/B, the linker sequence, Stem III, and the antisequestrator (nucleotides 1-166, **Fig. 1a**). However, it lacks the region spanning the ribosome binding site, thereby precluding the transition to the sequestrator conformation. During *in vitro* transcription, a DNA oligonucleotide complementary to the 3’ extension was added to the reaction. The DNA oligonucleotide carried the donor dye (Cy3) at its 3’ end and a biotin modification on the 5’ end. The labeled and biotinylated oligonucleotide allows for surface immobilization of the *Mtb-ileS* T-box via biotin-streptavidin interaction, and also places the donor dye in proximity to the 3’ terminus of the T-box construct for smFRET measurements (**Fig. 1c**) once hybridized (**Fig. 2a**). Positioning of the extension at the 3’ end of the riboswitch ensured that the decoding module would have an opportunity for vectorial folding during transcription, and that only full-length transcripts would anneal the oligonucleotide and be observed in smFRET imaging.

**Figure 2.**
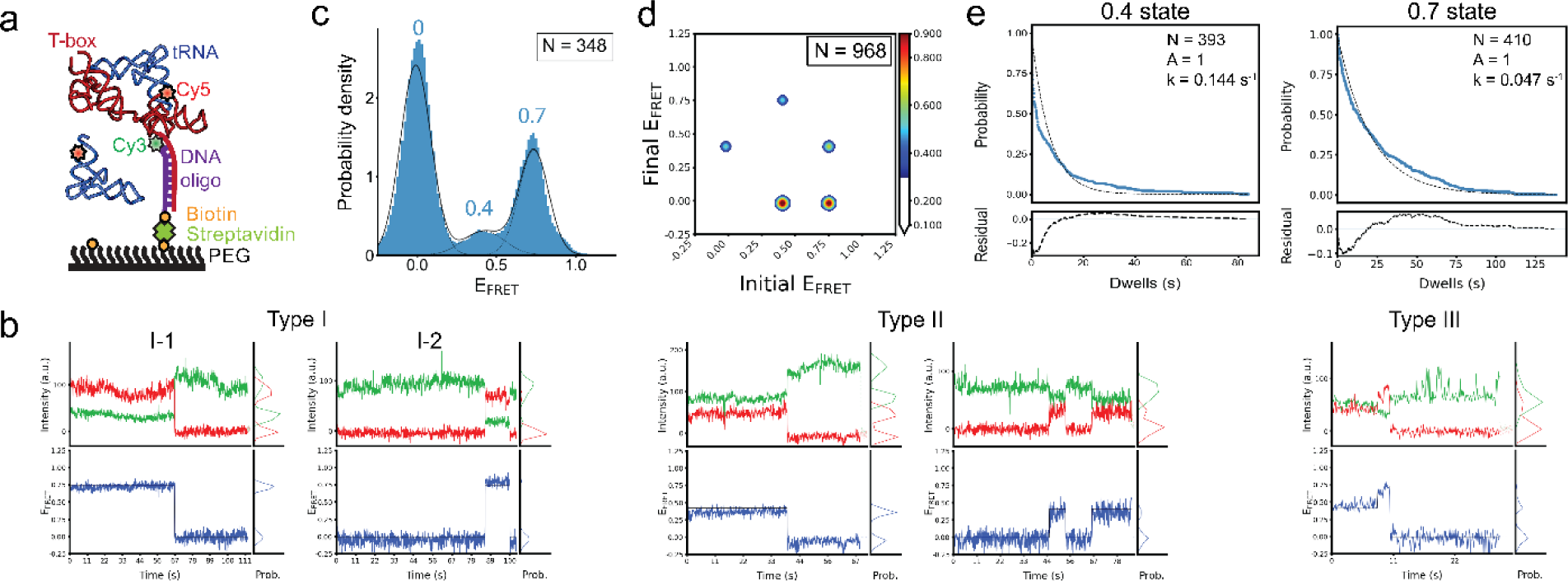
smFRET experiments show binding of tRNA^Ile^ to the wild-type *Mtb-ileS* T-box riboswitch. **a)** Schematic diagram illustrating the smFRET experiment. The T-box was anchored to the surface via hybridization to a DNA oligonucleotide (purple) biotinylated on the 5’ end and Cy3-labeled at the 3’ end. The tRNA^Ile^ was labeled on the 5’ end with Cy5. **b)** Representative Cy3 (green) and Cy5 (red) fluorescence intensity (top) and smFRET trajectories (bottom) of the *Mtb-ileS* T-box riboswitch/tRNA^Ile^. The smFRET efficiency was calculated as (I_Cy5_/(I_Cy3_-I_Cy5_).Three types of trajectories were observed: Type I trajectories only show FRET efficiency around 0.7, Type II trajectories only show FRET values around 0.4, and Type III trajectories sample the 0.4 and 0.7 states. **c)** FRET efficiency histogram of the data. Dotted black lines show the individual populations obtained from the consensus HMM modeling, while the model’s population-weighted set of efficiency distributions is plotted with a solid black line. N reports the number of traces included in the histogram. **d)** Transition density plot constructed using the idealized Viterbi paths modelled to the entire dataset using tMAVEN (26). The contour level colors report the normalized counts. N reports the total number of transitions. **e)** Dwell time survival plot for the 0.4 and 0.7 FRET states. The data were fitted using a single exponential decay function (Ae^(-tk)^). The number of events (N) used for the analysis and the fitting parameters (A,k) are shown. Data are shown in blue and the model in dotted black lines. The fit residuals are shown in the bottom plot.

The *Mtb ileS* T-box was purified directly from the transcription reaction by size exclusion chromatography (SEC) under conventional conditions (25). This process yielded readily distinguishable fractions of both unfolded and folded *Mtb-ileS* T-box states (**Supplementary** Fig. 1). The folded population of *Mtb-ileS* T-box molecules is consistent with their predicted apparent molecular weight, considering that the substantial negative charge of the molecule due to the phosphate backbone results in a larger hydration shell. The apparent molecular weight of an RNA is 3–5 times larger than a globular protein of comparable molecular weight and hence elutes from a SEC column significantly earlier (25). SEC purification also facilitated direct comparison of the overall folding of the wild-type (WT) construct with engineered mutants, ensuring that any binding differences were not due to misfolding (**Supplementary** Fig. 1). After transcription and SEC purification, the fraction corresponding to the fully transcribed and correctly folded *Mtb ileS* T-box riboswitch, already annealed to the biotinylated and Cy3 labeled DNA oligonucleotide, was used for smFRET imaging. In summary, the design with a 3’ extension, together with direct purification of the T-box construct from the transcription reaction via SEC avoided denaturation and refolding of the T-box riboswitch, which could disrupt its function. For intermolecular smFRET experiments with the cognate tRNA ligand, an acceptor fluorophore (Cy5) was conjugated at the 5’ end of the *M. tuberculosis* tRNA^Ile^ molecule (**Methods**). The purification of the tRNA was carried out by urea-PAGE, followed by refolding using standard procedures (**Methods**).

Biolayer Interferometry (BLI) binding assays were used to assess the effect of fluorescent labeling on tRNA binding to the Cy3-labelled *Mtb ileS* T-box (**Supplementary** Fig. 2a). For the BLI assays, the *Mtb ileS* T-box purified as described above was immobilized on the biosensor. tRNA association was measured in a solution containing uncharged tRNA^Ile^. After measuring the tRNA^Ile^ association, the biosensor was immersed in a tRNA^Ile^-free solution to enable tRNA^Ile^ dissociation detection. To fit the BLI data, three different binding models were considered: one-step binding, parallel one-step binding, and sequential two-step binding (**Supplementary** Fig. 2b). The binding and dissociation curves were fit most accurately by the two-step sequential binding model (**Supplementary** Fig. 2c). The apparent equilibrium dissociation constant (K_D_) was calculated using the two-step sequential binding model as described in the **Supplementary Information** section. The K_D_ of *Mtb ileS* T-box binding to tRNA^Ile^ obtained in these assays agrees well with previously reported affinities for this class of T-box (21). K_D_ with labeled tRNA^Ile^ (1.03 µM) was increased by only 2-fold compared to unlabeled tRNA^Ile^ (0.474 µM), demonstrating that the introduction of a label at the 5’ end of tRNA^Ile^ resulted in only a moderate impact on the binding affinity (**Supplementary** Fig. 2d**, Supplementary Table I**). This effect is milder than one which was previously reported for a similar experiment performed on the *B. subtilis glyQS* T-box riboswitch (19).

### Uncharged tRNA^Ile^ stably binds to the *Mtb ileS* T-box

Time-lapsed single-molecule fluorescence images of *Mtb ileS* T-box were captured in the presence of 100 nM tRNA^Ile^-Cy5 at equilibrium conditions (**Fig. 2b**). The distribution of FRET efficiency values resulting from tRNA^Ile^-Cy5 binding displays two main populations, with mean values of 0.4 and 0.7 (**Fig. 2c, Supplementary Table II**). Furthermore, the presence of a zero FRET state in the histograms can be attributed to both unbinding and acceptor fluorophore photobleaching events. The observed distribution is the result of three types of behaviors (**Fig. 2b and Supplementary Table III)**. Type I traces, comprising 77.3 % of the data, sample only the 0.7 FRET state in addition to the zero FRET state. Among Type I traces, the majority of traces (91.8 %) display a single, stable signal at 0.7 that undergoes single step photobleaching. A very small population (8.2 %) shows transitions from zero to 0.7 FRET efficiency. 17.8 % of traces (Type II) sampled only the 0.4 state in addition to the zero state, either once or several times before photobleaching. Finally, a very small fraction of traces (Type III traces, 4.9 %) sample both the 0.7 and 0.4 states.

Global Hidden Markov Modeling (HMM), wherein all trajectories are simultaneously modeled using a single HMM (26), further revealed the relative frequencies of transition events and associated transition rates (**Fig. 2d, Supplementary Tables IV** and **V**). Type I traces predominately demonstrate transitions from the 0.7 state to the zero FRET state (22.6%, calculated as the number of transitions of this type over the total number of transition events), with infrequent transitions from the zero to the 0.7 state (2.7 %) (**Supplementary** Fig. 3a). The minor Type II traces also contribute to a large number of transition events from 0.4 to zero FRET (25 %) and from zero to 0.4 FRET (12.8 %) (**Supplementary** Fig. 3b), as a fraction of Type II traces sample the 0.4 state multiple times in each trajectory. 0.4 to zero FRET transitions are more frequent than the reverse transitions as the former also include the photobleaching step. Finally, Type III traces demonstrate inter-state transitions between 0.7 and 0.4 FRET states (20.1% for 0.7 to 0.4 transitions and 16.8 % for 0.4 to 0.7 transitions) (**Supplementary** Fig. 3c).

According to the crystal structure (9), there should be a distance of approximately 40 Å between the labeling positions at the 3’ end of the *Mtb ileS* T-box and the 5’ end of the tRNA (**Fig. 1c**). The 0.7 FRET value obtained from the measurements is within the predicted range of FRET values, assuming a Förster distance of 40 - 50 Å for Cy3-Cy5 FRET pair (27). Hence, it is plausible to designate the 0.7 FRET state as the fully bound state of tRNA^Ile^ to the *Mtb ileS* T-box. The large population of Type I traces and 0.7 to zero transitions indicate that, preponderantly, the tRNA^Ile^ was fully bound to the T-box prior to data collection. This state persisted until the acceptor fluorophore photobleached, thus restricting the observed lifetime of the 0.7 state to around 21.1 s (**Fig. 2e** and **Supplementary Table VI**). The characteristics of the 0.7 state are strikingly similar to those of the fully bound state of *B. subtilis glyQS* T-box riboswitch in complex with uncharged tRNA^Gly^ (19). This similarity is expected due to the structural conservation between antisequestrator and antiterminator domains of translational and transcriptional T-boxes, respectively, in complex with their cognate tRNAs (**Supplementary** Fig. 3d) (9,15).

While the nature of the 0.4 FRET state is not clear from the data above, there are more frequent transitions from the zero to the 0.4 state than from the zero to the 0.7 state, suggesting that it is more likely for tRNA^Ile^ to bind this unknown intermediate conformation associated with the 0.4 state than the fully bound state. (**Fig. 2d, Supplementary** Figs. 3a and b**, Supplementary Table IV**). It should be noted that the binding rate of tRNA^Ile^ to either the unknown intermediate or fully bound states from the unbound state is likely to be significantly underestimated by these experiments, given that the majority of the population probed was already in the fully bound state during the data acquisition time window.

### Anticodon recognition leads to a partially bound state of the tRNA

The nature of the 0.4 state cannot be deduced from the available structural data. However, it resembles a state observed in studies of the *B. subtilis glyQS* T-box riboswitch (19). In the *glyQS* T-box, a 0.4 state corresponds to a partially bound state in which only the anticodon/specifier interactions have been established. To test whether the 0.4 FRET state in *Mtb ileS* T-box represents such partially bound state, two experiments were devised. In the first experiment, smFRET measurements were conducted with the *Mtb ileS* T-box using a tRNA^Ile^ variant lacking the terminal NCCA-3’ sequence (tRNA^Ile-ΔNCCA^-Cy5). It was anticipated that tRNA^Ile-ΔNCCA^-Cy5 would only be able to bind to the decoding module but not to the discriminator domain (**Fig. 3a**). With the use of this tRNA substrate, two apparent populations were initially identified in the FRET efficiency distribution: a 0.5 FRET state and a zero FRET state (**Fig. 3b, Supplementary Table II**). The largely diminished 0.7 FRET state further supports the assignment of the 0.7 FRET to represent the fully bound state. Hence, the presence of the 0.5 FRET efficiency state was ascribed to a partially bound state in which no interactions with the NCCA sequence had occurred and only the tRNA anticodon region was bound. Global HMM (26) was used to infer transition frequencies and rates. This analysis revealed that the 0.5 FRET state is sampled more transiently compared to the 0.7 FRET state observed with tRNA^Ile^-Cy5 (**Fig. 3c, Supplementary Table IV**). The transition rate from the zero state to the 0.5 state was 0.019 s^-1^, while the transition rate from the 0.5 state to the zero state was 0.0532 s^-1^ (**Supplementary Table V**), corresponding to an average lifetime of 18.8 seconds before tRNA dissociation (**Supplementary Table VI**).

**Figure 3.**
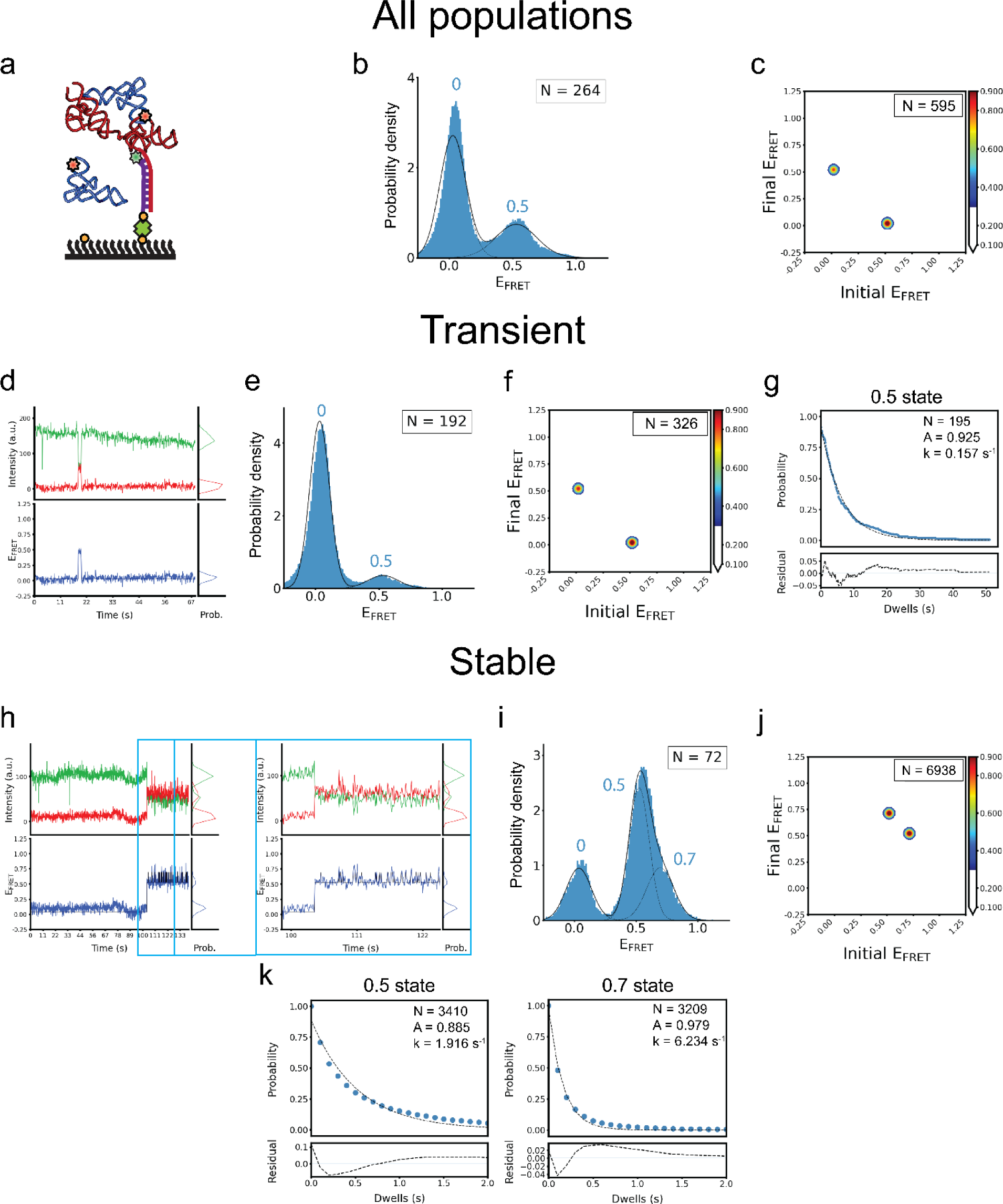
smFRET experiments with the tRNA^Ile-ΔNCCA^ mutant show consequences of the absence of the NCCA/specifier interactions. **a)** Schematic diagram illustrating the smFRET experiment with the tRNA^Ile-ΔNCCA^-Cy5 construct. **b)** FRET efficiency histogram for tRNA^Ile-ΔNCCA^-Cy5 mutant. Fitted populations using consensus HMM modeling are shown as in Figure 2c. N reports the number of traces in the histogram. **c)** Transition density plot for tRNA^Ile-ΔNCCA^-Cy5 mutant for all data constructed using the idealized Viterbi paths modelled with tMAVEN (26), plotted the same way as described in Figure 2d. **d)** Representative Cy3 (green) and Cy5 (red) fluorescence intensity (top) and smFRET trajectories (blue, bottom) for the subpopulation of the *Mtb-ileS* T-box riboswitch/ tRNA^Ile-ΔNCCA^-Cy5 complex displaying transient bindings. **e)** FRET efficiency histogram of the subpopulation of tRNA^Ile-ΔNCCA^-Cy5 mutant displaying transient binding. **f)** Transition density plot for the transient traces constructed using the idealized Viterbi paths for the subpopulation displaying transient bindings. **g)** Dwell time survival plots for the 0.5 state in the transient binding subpopulation. The data were fitted using a single exponential decay function (Ae^(-tk)^). The number of events (N) used for the analysis and the fitting parameters (A,k) are shown. Data and fitting results as described in Figure 2e. **h)** Representative fluorescence intensity and smFRET trajectory from the subpopulation of the *Mtb-ileS* T-box riboswitch/ tRNA^Ile-ΔNCCA^-Cy5 complex displaying stable binding (Left). A zoom of a region framed by cyan boxes is displayed on the right, which displays rapid transitions between the 0.5 and 0.7 states. **i)** FRET efficiency histogram of the subpopulation of tRNA^Ile-ΔNCCA^-Cy5 mutant displaying stable binding. **j)** Transition density plot constructed using the idealized Viterbi paths for the stable binding subpopulation. **k)** Dwell time survival plots for the 0.5 and 0.7 states in the stable traces and fitting result using a single exponential decay function. Data and fitting results are presented as described in Figure 2e.

In a second experiment, a mutant of the *Mtb ileS* T-box which prevents tRNA^Ile^ interactions through the NCCA sequence was constructed by eliminating the discriminator domain (Δ-Discriminator Mutant, nucleotides 1-93). The SEC elution profile of the Δ-Discriminator Mutant resembles that of the wild-type *Mtb ileS* T-box, with the exception that the peak of interest eluted at a later volume (**Supplementary** Fig. 1). This shift was anticipated due to the lower molecular weight of the construct. The deletion of the discriminator domain had an impact on the ability of the T-box to bind to tRNA^Ile^, as demonstrated by BLI measurements. The K_D_ increased from 0.474 µM for the wild type, to 4.55 µM for the mutant (**Supplementary Table I**), corresponding to a ∼10-fold reduction in the binding affinity. smFRET analysis on this mutant using tRNA^Ile^-Cy5 (**Fig. 4a**) yielded two distinct populations centered at zero and 0.66 FRET respectively (**Fig. 4b,c**, **Supplementary Table II**). Global HMM enabled quantification of inter-transitions between the zero and 0.66 FRET states (**Fig. 4d, Supplementary** Fig. 4g). The transition rate from the zero to the 0.66 state was 0.0134 s^-1^, whereas the transition rate from the 0.66 state to the zero state was 0.0568 s^-1^, corresponding to an average lifetime of 17.6 seconds for the 0.66 FRET state (**Fig 4e**), resembling the lifetime of the 0.5 FRET state in the case of WT T-box with tRNA^Ile-ΔNCCA^ (**Supplementary Tables V** and **VI**).

**Figure 4.**
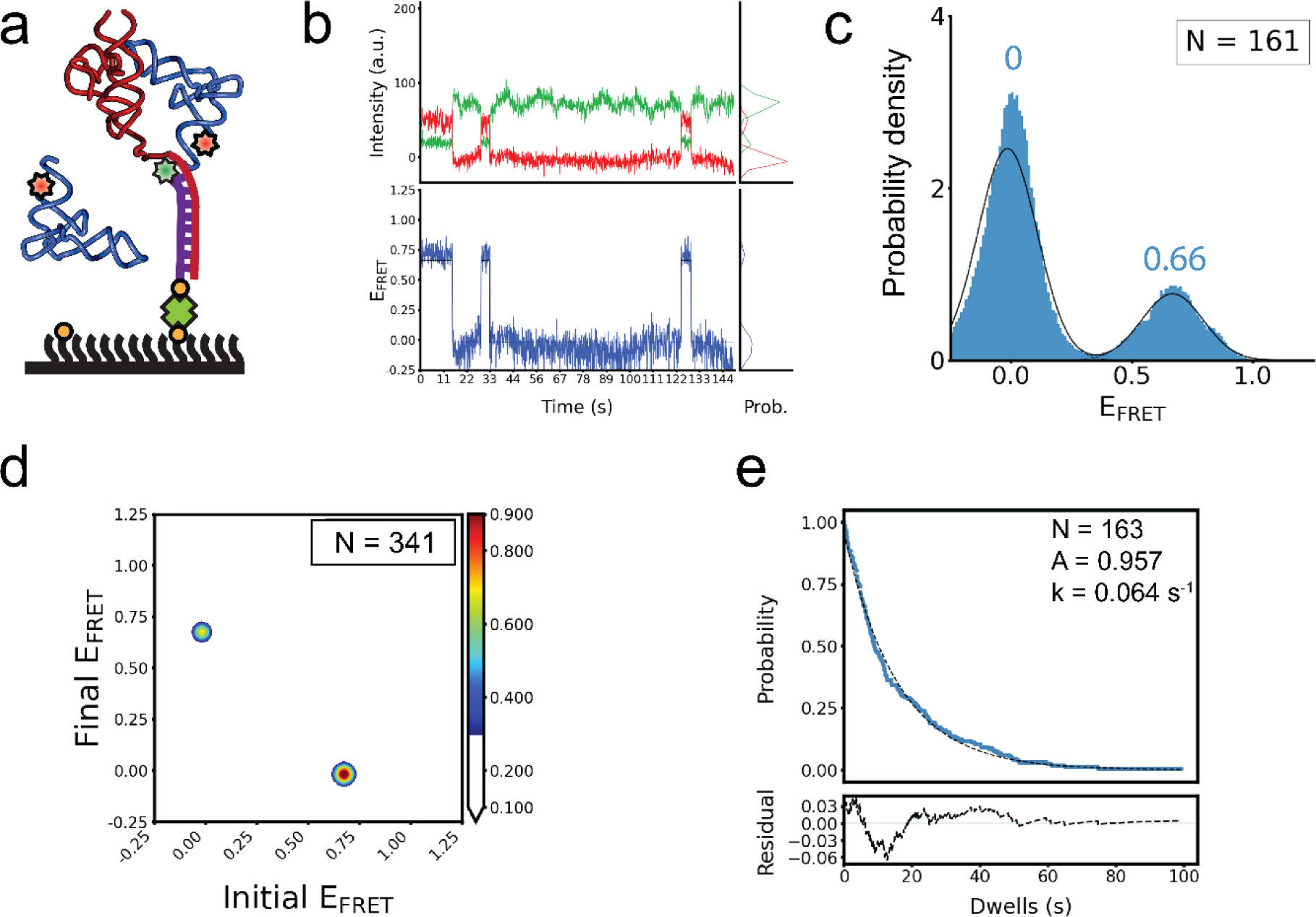
smFRET experiments using the Δ-Discriminator Mutant. **a)** Schematic diagram illustrating the smFRET experiment with the *Mtb-ileS* T-box riboswitch Δ-Discriminator Mutant and tRNA^Ile^. **b)** Representative Cy3 (green) and Cy5 (red) fluorescence intensity (top) and smFRET trajectories (blue, bottom) of the *Mtb-ileS* T-box riboswitch Δ-Discriminator Mutant and tRNA^Ile^ complex. **c)** FRET efficiency histogram of the data with the Δ-Discriminator Mutant. Fitted populations using consensus HMM modeling are plotted as in Figure 2c. N reports the number of traces in the histogram. **d)** Transition density plot constructed using the idealized Viterbi paths, as described in Figure 2d. N reports number of transitions in the plot. **e)** Dwell time survival plots and fitting results using a single exponential decay function (Ae^(-tk)^). The number of events (N) and the fitting parameters (A,k) are shown. Data are shown in blue and the model in dotted black lines. The fit residuals are shown in the bottom plot. Data and fitting results are presented as described in Figure 2e.

For the Δ-Discriminator Mutant, the estimated distance between the truncation position and the 5’ end of the tRNA is around 29 Å (**Fig. 1c**). The observed higher FRET efficiency at 0.66 is to be expected with a Förster distance of 40 to 50 Å (27). This effect is likely caused by a collapse of the single-stranded linker region in the absence of the discriminator domain, with which it normally establishes contacts (**Fig. 1c**). However, as the transition rates between the zero and 0.5 states in the wild-type + tRNA^Ile-ΔNCCA^-Cy5 and the zero and 0.66 states in the Δ-Discriminator mutant are nearly identical (**Supplementary Table V**), and, as in both cases the NCCA-discriminator interactions are perturbed, both the 0.5 and the 0.66 FRET states were interpreted to correspond to the same partially bound state. The lifetimes of the partially bound states in these two cases were two to three times longer compared to the lifetime of the 0.4 FRET state observed in the case of the WT *Mtb ileS* T-box with tRNA^Ile^ (**Supplementary Table VI**). This difference is expected however, given the fact that the 0.4 FRET state in WT *Mtb ileS* T-box with tRNA^Ile^ is capable of transitioning to the fully bound state. The absence of the NCCA sequence may allow the decoding module to shift the 5’ end of tRNA^Ile-ΔNCCA^-Cy5 closer to the discriminator domain, leading to an apparently higher FRET efficiency state compared to the 0.4 state. Taken together, the results presented so far suggest a two-step binding model in which the interactions between the tRNA NCCA and anticodon with the T-box do not happen simultaneously.

### tRNA anticodon binding precedes NCCA recognition in *Mtb ileS* T-box

The above experiments demonstrate that the *Mtb ileS* T-box can identify the anticodon region of tRNA^Ile^ even in the absence of the NCCA sequence. Furthermore, two previous studies with translational T-boxes missing the discriminator domain (23,24) corroborate this finding. Nevertheless, to determine if interactions between *Mtb ileS* T-box and the NCCA sequence are possible without anticodon recognition, additional experiments were pursued. First, tRNA^Trp^-Cy5 was introduced into a flow chamber containing pre-immobilized *Mtb ileS* T-box, but no binding of tRNA^Trp^-Cy5 to the *Mtb ileS* T-box was detected. Binding by BLI was also not detectable. Second, an additional T-box construct was designed in which the specifier region, responsible for identifying the cognate anticodon, was mutated to recognize tRNA^Trp^ instead (AUC mutated to UGG), hereafter referred to as the Specifier Mutant. The Specifier Mutant fraction eluted slightly earlier than the WT *Mtb ileS* T-box in SEC analysis, likely due to a potential contribution of the specifier sequence in maintaining the native conformational landscape of the decoding module (24). No binding was observed when tRNA^Ile^-Cy5 was added to a flow-chamber pre-immobilized with the Specifier Mutant. Consistent with the smFRET measurements, BLI assays measured the K_D_ of tRNA^Ile^ binding to the Specifier Mutant to be 160 µM, three orders of magnitude larger than the K_D_ of WT *Mtb ileS* T-box (0.474 µM) (**Supplementary Table I**), and well above the expected physiological concentrations of tRNA (28).

Overall, the results of these experiments confirm that tRNA binding requires the specifier region to recognize the tRNA anticodon before interactions with the NCCA sequence can be established. The experiments also suggest that the translation-regulating *Mtb ileS* T-box also follows the same sequential binding mechanism as the transcription-regulating *B. subtilis glyQS*

T-box.

### The partially bound state in *Mtb ileS* T-box exhibits conformational heterogeneity

While *Mtb ileS* T-box follows the same sequential binding mechanism as *B. subtilis glyQS* T-box, the presence of conformational heterogeneity in the partially bound state of *Mtb ileS* T-box complex with tRNA^Ile-ΔNCCA^-Cy5 was observed. This conformational heterogeneity is absent in the *B. subtilis glyQS* T-box case (19). Specifically, two distinct types of traces were observed upon tRNA^Ile-ΔNCCA^-Cy5 binding to the WT *Mtb ileS* T-box: a population where the 0.5 state is sampled transiently (72.7 %), and a second population where the 0.5 FRET state is stable until photobleaching (27.3 %) (**Fig. 3d, h** and **Supplementary Tables II** and **III**). The presence of conformational heterogeneity was also recognized in the dwell time analysis of the 0.5 FRET state, which cannot be described by using a single exponential decay function (**Supplementary** Fig. 4b).

In light of this finding, the stable and transient subpopulations were further analyzed separately. For the subpopulation that transiently sampled the 0.5 state (**Fig. 3d, e**), transitions were identified via composite HMM, i.e., when individual trajectories are modeled separately (26), which is more sensitive to kinetic heterogeneity. Analysis of transitions show homogeneous sampling of the 0.5 state in both directions (**Fig. 3f, Supplementary** Fig. 4c) The dwell time of the 0.5 FRET state in this subpopulation can be fit with a single-exponential decay function, yielding an average lifetime of 6.3 s (**Fig. 3g** and **Supplementary Table VI**). This subpopulation was reminiscent of the case of tRNA^Gly-ΔNCCA^-Cy5 binding to the WT *B. subtilis glyQS* T-box, in which only transient sampling of the partially bound state was observed, and the dwell time of the partially bound state was described well with single exponential decay function with a similar lifetime (19).

Remarkably, molecules that spent a longer time in the 0.5 FRET state actually displayed rapid transitions to a 0.7 FRET state, the latter reminiscent of the FRET efficiency observed in the fully bound state (**Fig. 3h, i**). In this subpopulation, the 0.7 FRET state was very transiently sampled with 1-2 time points (< 200 ms), which was not captured by the global HMM algorithm. In order to model accurately this fast transition behavior, Gaussian mixture modelling (GMM) (26) was used (see **Methods**). Since the short-lived 0.7 FRET state is consistently sampled across the dataset of this subpopulation, this approach, which assumes that all the data points originate from a mixture of a finite number of Gaussian distributions with unknown parameters, is capable of detecting it. The GMM analysis revealed that transition events occurred predominantly between the 0.7 and 0.5 FRET states (92.2 % of total transitions) (**Fig. 3j** and **Supplementary Table IV**). These FRET states displayed short lifetimes, 0.52 s and 0.16 s for the 0.5 and 0.7 FRET states, respectively (**Fig. 3k**). These results suggest that, upon anticodon recognition, the *Mtb ileS* T-box exhibits fast conformational dynamics, unlike the *B. subtilis glyQS* T-box. In the absence of the 3’ terminal NCCA sequence in tRNA^Ile^, some *Mtb ileS* T-box molecules undergo futile attempts to achieve the fully bound state. In other words, the fully bound state can be transiently sampled without establishment of the NCCA-discriminator interaction. However, establishment of the NCCA-discriminator interaction in the presence of uncharged tRNA^Ile^ stabilizes the fully bound state by at least 130-fold.

### tRNA binding alters the conformational equilibrium of *Mtb ileS* T-box

To study intramolecular conformational changes of the *Mtb ileS* T-box upon tRNA binding, a new construct was designed, termed IntraFRET. This construct has a 5’ extension that enables it to bind a 5’-end Cy5-labeled DNA oligonucleotide while retaining the 3’ extension used for immobilization and Cy3 labeling (**Fig. 5a**). This construct enables probing the distance changes between the 5’ and 3’ ends of the *Mtb ileS* T-box. The SEC elution profile of the IntraFRET molecule exhibited greater complexity as compared to the *Mtb ileS* T-box construct with only the 3’ oligonucleotide attached. This was evident by the appearance of several peaks in the chromatogram (**Supplementary** Fig. 1). By measuring both Cy3 and Cy5 absorbance signals from each SEC fraction, only one of the peaks was found to contain both signals, and this fraction was used for further smFRET analysis. This peak, corresponding to the doubly labeled T-box, eluted earlier compared to T-box with only the 3’ oligonucleotide due to its higher molecular weight.

**Figure 5.**
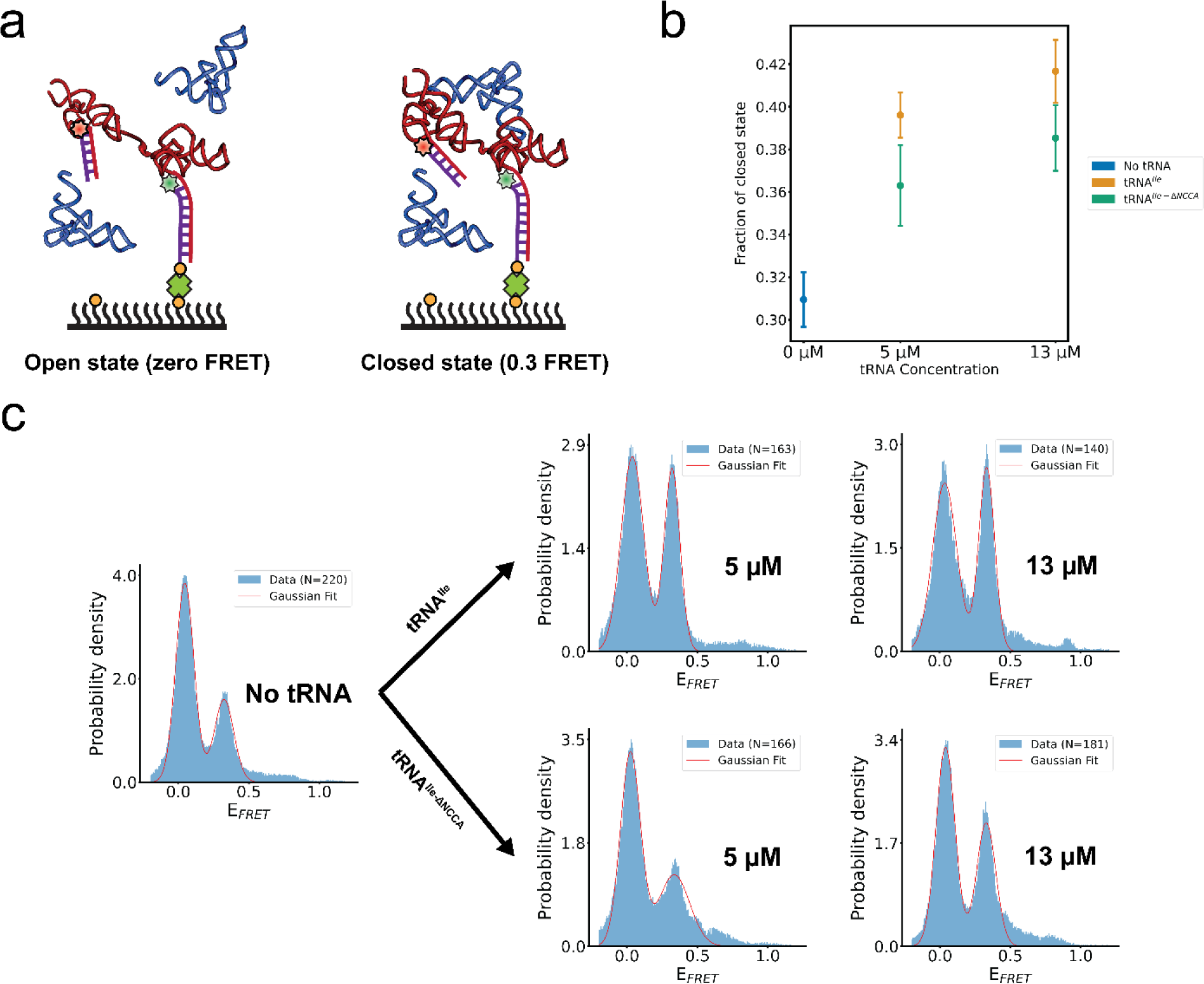
The *Mtb-ileS* T-box riboswitch samples open and closed conformations. **a)** Schematic diagram illustrating the smFRET experiment using an intramolecularly labeled *Mtb-ileS* T-box construct. Two FRET efficiency states were observed, assigned to an open (zero FRET, left) and a closed (0.3 FRET, right) state. **b)** FRET efficiency histograms of intramolecularly labeled *Mtb-ileS* T-box in the absence of tRNA (left), and in the presence of different concentrations of tRNA^Ile^ (top) and tRNA^Ile-ΔNCCA^ (bottom). The orange line corresponds to the fits of a Gaussian function to each peak. N reports the number of traces in each histogram. Only the first 40 frames from FRET trajectories are included in the histogram to avoid including data after photobleaching of the fluorophores. **c)** Changes in the fraction of the closed conformation as a function of tRNA^Ile^ or tRNA^Ile-ΔNCCA^ concentration. The fraction of the closed state was calculated by the integrated peak area corresponding to the 0.3 FRET state divided by the sum of the peak areas corresponding to the zero and the 0.3 FRET states in **b**).

For smFRET measurements, the IntraFRET molecules were immobilized to the imaging slide and time-lapsed fluorescence images were recorded with and without tRNA. To distinguish between molecules in a potential zero FRET state from those with only a Cy3 label, five frames were recorded using direct Cy5 excitation before time-lapsed FRET imaging. In the absence of tRNA, the IntraFRET sample showed two distinct FRET states centered at zero and 0.3, respectively (**Fig. 5b**). We assigned these states as open (zero FRET) and closed (0.3 FRET). The closed state is consistent with the expected distance of around 68 Å between the 3’ and 5’ ends of the T-box in complex with tRNA^Ile^ (**Fig. 1c**) (27). Addition of tRNA^Ile^ caused a dose-dependent increase in the proportion of the closed state (**Figures 5b, c**). Remarkably, addition of tRNA^Ile-ΔNCCA^ similarly increased the proportion of the 0.3 FRET efficiency state in a dose-dependent manner, albeit with a reduced efficiency (**Figs. 5 b, c**). These findings suggest that in the absence of the tRNA ligand, the *Mtb ileS* T-box samples both the open and closed conformations autonomously. The experiments with tRNA^Ile-ΔNCCA^ indicate that anticodon recognition alone increases the population of the closed state. Interactions between the NCCA end of the tRNA and the discriminator of the T-box further enhance the stability of the closed state. These observations are in stark contrast to equivalent experiments with the *glyQS* T-box, where an intramolecular conformational change is exclusively associated with the binding of both the anticodon and NCCA (19).

### Stem IIA/B-linker interactions allosterically modulate tRNA binding

The linker region in *Mtb ileS T-box* (nucleotides 79–92) connects the decoding module and the discriminator domain (**Fig. 1a**). This linker region is predominantly stabilized by extensive interactions with Stem IIA/B, resulting in limited flexibility. Specifically, nucleotide A81 at the 5’ end of the linker forms an A-minor interaction with the G68-C77 pair in Stem IIA/B. Then, the 5′-half of the linker rests against the interface between Stems II and IIA/B and forms a pseudoknot structure by base pairing with three of the nucleotides in the loop of Stem IIA/B (**Fig. 1a**). Hence, it has been suggested that this pseudoknot imposes the constraints required to position the discriminator domain in proximity to the tRNA acceptor arm (9). Disruption of the aforementioned A-minor interaction (A81G) was shown to significantly decrease the functional efficiency of the *B. subtilis tyrS* T-box (9). To investigate how the Stem II-A/B-linker interactions modulate the tRNA binding kinetics, an *Mtb ileS* T-box mutant that included the A81G substitution as well as mutations expected to disrupt the three critical base pairs with Stem IIA/B was generated (**Fig. 6a**). This mutant is hereafter referred to as the “Linker Mutant”. The SEC elution profile and K_D_ (measured by BLI) of the Linker Mutant were similar to those of the WT *Mtb ileS* T-box (**Supplementary** Figs. 1, 2b**, Supplementary Table I**).

**Figure 6.**
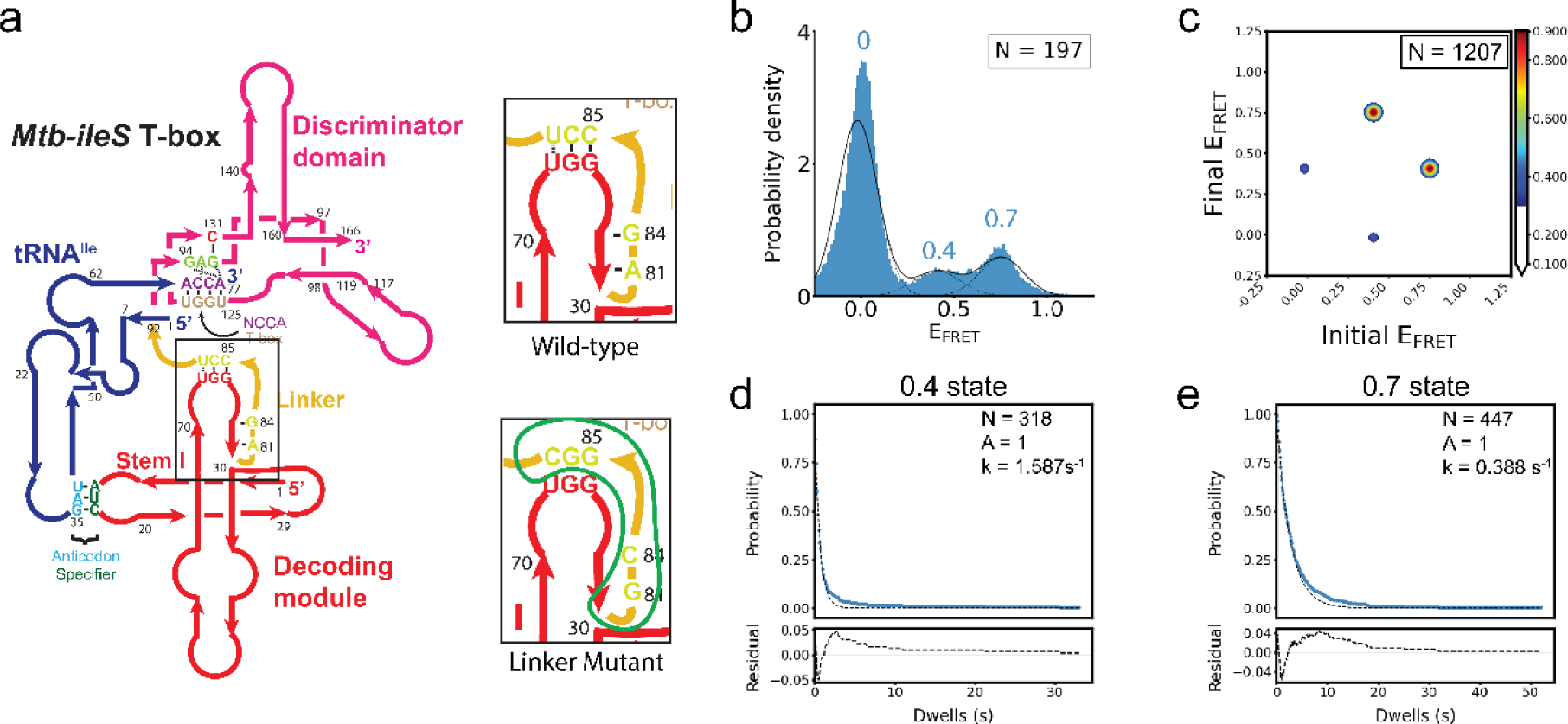
Linker Mutations destabilize the *Mtb-ileS* T-box/tRNA complex. **a) Left**, schematic diagram of the *Mtb-ileS* T-box with the linker region shown inside the black box. **Right,** schematic diagrams showing the linker region (top) and the mutations introduced in the Linker Mutant (bottom). The region affected is circled in green in the mutant diagram. **b)** FRET efficiency histogram of the data with the Linker Mutant. Fittedpopulations using consensus HMM modeling are plotted as in Figure 2c. N reports the number of traces in the histogram. **c)** Transition density plot constructed using the idealized Viterbi paths, as described in Figure 2d. N reports number of transitions in the plot. Dwell time survival plots and fitting result using a single exponential decay function (Ae^(-tk)^) for the 0.4 FRET state **(d)** and 0.7 FRET state **(e)**. The number of events (N) and the fitting parameters (A,k) are shown. Data are shown in blue and the model in dotted black lines. The fit residuals are shown in the bottom plot. Data and fitting results are presented as described in Figure 2e.

The FRET values resulting from the binding of tRNA^Ile^-Cy5 to the Linker Mutant form two populations, with mean values of 0.4 and 0.7 (**Fig. 6b, Supplementary Table II**), similar to the ones observed using the WT *Mtb ileS* T-box. However, the 0.7 FRET state is much less populated in this mutant compared to the WT T-box. With the same categorization as the one utilized for the WT construct (**Supplementary** Fig. 5a), Type I, II, and III traces comprise 20.3%, 46.19%, and 33.5% of the total traces, respectively (**Supplementary Table III**). The reduced fraction of Type I traces, and the corresponding increase in the fractions of Type II and III traces suggest that the stability of the fully bound state is compromised in this T-box mutant.

The transitions between 0.7 and 0.4 FRET states in both directions are most frequent, accounting for 63.3% of all transitions (**Fig. 6c** and **Supplementary Table IV**). The transition rates between the 0.7 and 0.4 states are 8- to 15-fold faster in the Linker Mutant compared to those in the WT construct (**Supplementary Table V)**. The transitions between the 0.4 and the zero FRET states contribute to 29.4% of all transitions (**Fig. 6c, Supplementary** Fig. 5b and **Supplementary Table IV**), with the rates around 2- to 4-fold faster compared to the WT construct (**Supplementary Table V)**. Similar to the WT construct, transitions between zero and 0.7 FRET states were the least frequent, accounting for only 7.3% of the total (**Fig. 6c, Supplementary** Fig. 5b and **Supplementary Table IV**). The average lifetime of the 0.4 state was 630 ms, a ∼10-fold decrease compared to the WT construct (**Figs. 2e, 6d** and **Supplementary Table VI**). Consistent with increased transition rates between the states, the mean lifetime of the 0.7 state was 2.6 seconds (**Fig. 6e** and **Supplementary Table VI**), which also representing a ∼10-fold decrease compared to the WT *Mtb ileS* T-box (**Fig. 2e**). These results demonstrate the importance of the A-minor interaction and pseudoknot structure in maintaining the stability of both the partially and fully bound states. Overall, these results confirm the significance of the *Mtb ileS* T-box linker region in tRNA binding and point to allosteric effects on tRNA recognition, since the Stem II A/B-linker interactions are not directly involved in tRNA binding.

### The conserved RAG sequence confers specificity and stability to NCCA-Discriminator interactions

Prior structural data have provided insights into aminoacylation state sensing by the T-box discriminator domain. Specifically, a conserved consensus sequence RAG (R = purine) plays a crucial role in this process (9). In the *Mtb-ileS* T-box, the RAG sequence starts with G94, which interacts with an adenosine (A77) at the 3’ of the tRNA and with an antisequestrator ribose (U125). The second nucleotide in the sequence, A95, is crucial in facilitating continuous stacking between T-box nucleotides A129 and C131. Lastly, G96 binds C131 in the antisequestrator (9). These interactions provide the specificity required to bind an uncharged tRNA containing a terminal adenosine with intact 2′ and 3′ hydroxyl groups. This high level of specificity renders the NCCA binding pocket incompatible with any other chemical moieties, such as an attached amino acid. Indeed, single nucleotide substitutions in the RAG sequence (R94U, A95G, and G96C) significantly decrease the induction of the *B. subtilis tyrS* T-box (9). To investigate how this RAG sequence contributes kinetically to NCCA recognition, a mutant, termed “RAG mutant,” containing substitutions R94C, A95G, and G96C (**Fig. 7a**) was designed. The RAG mutant has a SEC elution profile that closely resembles that of the WT construct (**Supplementary** Fig. 1) and exhibits a moderate decrease in binding affinity to tRNA^Ile^, with a K_D_ of 1.48 µM, compared to 0.474 µM in the WT construct (**Supplementary Table I**).

**Figure 7.**
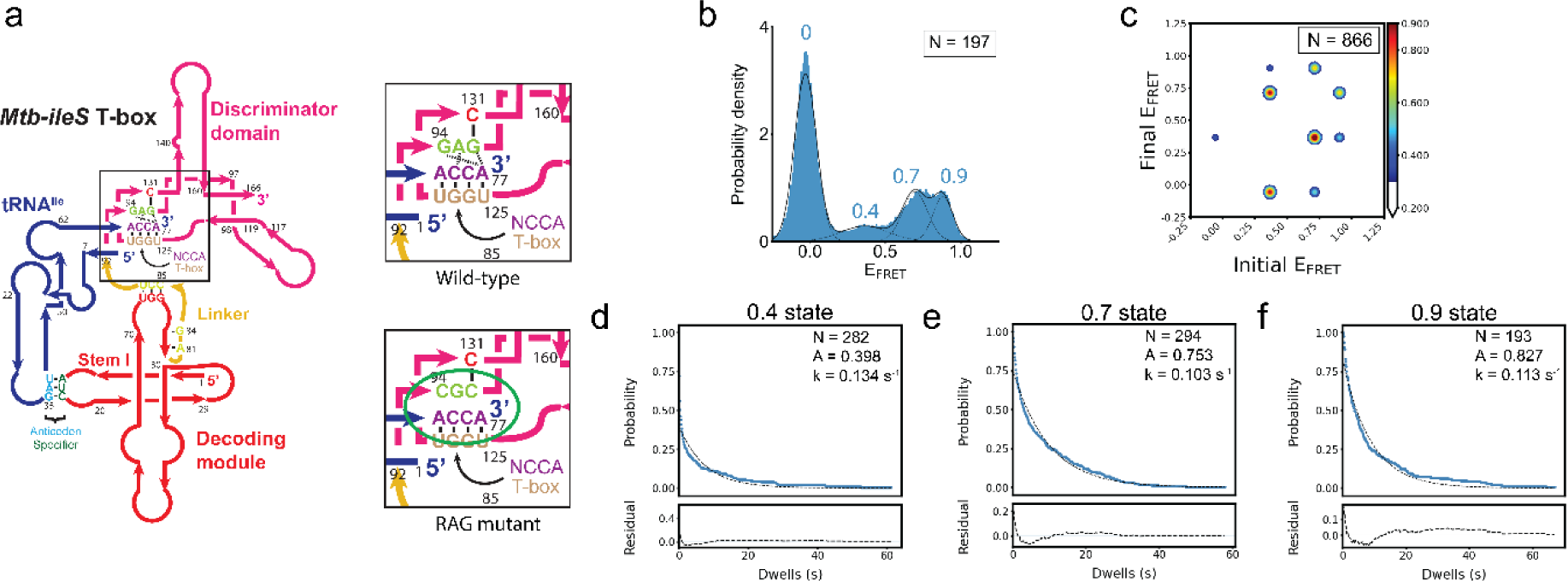
The RAG sequence is important for aminoacylation state recognition. a) Left: Schematic diagram of the *Mtb-ileS* T-box with the NCAA/T-box and RAG sequence regions shown inside the black box. **Right,** schematic diagrams showing the RAG sequence region (GAG) (top) and the mutations introduced in the RAG Sequence mutant (bottom). The interactions affected are circled in green in the mutant diagram. **b)** FRET efficiency histogram of the data with the Linker Mutant. Fitted populations using consensus HMM modeling are plotted as in Figure 2c. N reports the number of traces in the histogram. **c)** Transition density plot constructed using the idealized Viterbi paths, as described in Figure 2d. N reports number of transitions in the plot. Dwell time survival plots and fitting result using a single exponential decay function (Ae^(-tk)^) for the **d)** 0.4 FRET state, **e)** 0.7 FRET state and the **f)** 0.9 FRET state. The number of events (N) and the fitting parameters (A,k) are shown. Data are shown in blue and the model in dotted black lines. The fit residuals are shown in the bottom plot. Data and fitting results are presented as described in Figure 2e.

Interestingly, smFRET measurements using the RAG mutant and tRNA^Ile^-Cy5 revealed a new FRET state centered at 0.9, in addition to the 0.4 and 0.7 states (**Fig. 7b, Supplementary Table II**). Approximately 82.2% of the traces (Type I, II and IV) probe only one FRET state (0.7, 0.4, or 0.9 respectively) either once or a few times before photobleaching, while the remaining 17.8% of traces exhibit transitions between two or three states collectively (**Supplementary** Fig. 6a). Specifically, the Type II traces, representing traces only sampling of the 0.4 state, represent 17.8% of the data, similar to the WT construct. Traces sampling only the 0.7 (Type I) and only the 0.9 (Type IV) FRET states add up to 64.4%, close to the fraction of the Type I traces in the WT T-box (**Supplementary** Fig. 6a, b and **Supplementary Table III**). This suggests that the fully bound state in the RAG mutant has two conformations corresponding to the 0.7 and 0.9 FRET states respectively. While the 0.7 FRET state corresponds to the same conformation of fully bound state observed in the WT construct, the 0.9 FRET state corresponds a new state only observed in the RAG mutant complex, hereafter referred to as the pseudo fully bound state.

The RAG mutant showed a similar frequency of transitions between 0.7 and 0.4 FRET states compared to the WT complex, albeit with two to three times faster transition rates (**Supplementary Tables IV** and **V**). In addition, transitions also occurred between the 0.7 and 0.9 FRET states with similar frequency as the 0.7 and 0.4 transitions, suggesting that the RAG mutant complex exhibits additional conformational dynamics between the fully bound state and the pseudo fully bound state. Finally, direct transitions between 0.4 and the 0.9 FRET states were also observed with less frequency **(Fig. 7c** and **Supplementary Table IV)**. Kinetic analysis determined the average lifetime of the 0.4 state to be 7.5 seconds (**Fig. 7d** and **Supplementary Table VI**), similar to that of the WT construct (**Fig. 2e** and **Supplementary Table VI**). However, the average lifetime of the 0.7 state is 9.7 seconds (**Fig. 7e**), reduced by approximately two-fold compared to the lifetime of the 0.7 state in WT T-box. The lifetime of the 0.9 state is 8.8 seconds (**Fig. 7f** and **Supplementary Table VI**), comparable to the lifetime of the 0.7 state for this mutant. The lifetime comparison between the RAG mutant and WT construct for each FRET state indicates that: 1) the fully bound states still represent a more energetically stable conformation than the partially bound state, and 2) the mutations make the NCCA-discriminator interactions less structurally constrained by generating an energetically similar pseudo fully bound state. These findings support the notion that the conserved RAG sequence plays a crucial role in regulating the ability of the discriminator domain to sense aminoacylation, likely by restricting potential degrees of freedom of the bound ligand.

### Stem II integrity is essential for proper folding of *Mtb ileS* T-box

In an attempt to investigate its dynamics and involvement in tRNA^Ile^ binding, Stem II was systematically shortened to generate three additional constructs, termed Stem II mutants A, B, and C (**Supplementary** Figure 7). Amongst the three, only Stem II mutant A produced a distinct peak in the SEC chromatogram (**Supplementary** Fig. 1). These findings suggest that Stem II mutants B and C are likely to fold into defective conformations. Stem II mutant A did not exhibit any measurable tRNA^Ile^ binding, as measured by either smFRET or BLI, hence precluding further study. These results emphasize the importance of the integrity of the decoding module in binding tRNA^Ile^. It was shown in recent single molecule and Small Angle X-ray Scattering (SAXS) studies of the decoding module that the formation of the anticodon binding groove depends on the conformational dynamics between Stems I, II, and the Stem IIA/B pseudoknot (23,24). The mutants presented hereby support that preserving the integrity of all stems is essential for the decoding module to maintain its natural dynamic ensemble. These results highlight the cooperative nature of the conformational landscape of *Mtb ileS* T-box.

## Discussion

There has been a surge in interest in the conformational dynamics and tRNA ligand binding mechanisms of translational T-box riboswitches. The folding landscape of the decoding module (containing Stems I, II, and IIA/B) of the *Nocardia farcinica ileS* T-box riboswitch and its coupling to tRNA binding have been explored in recent studies (23,24). Collectively, these studies showed that the decoding module of a translation-regulating T-box riboswitch exhibits a wide distribution of conformations, which undergo slow interconversion. tRNA binds directly to all conformers, each with unique kinetics. Binding promptly induces rapid rearrangement of the structural elements forming the decoding module, leading to a final, binding competent conformation. Furthermore, following dissociation, tRNA facilitates the return of the decoding module to its original conformation prior to binding (24). In addition, it has been shown that Mg^2+^ plays an important role in the global folding of the decoding module (23). Here, using a larger construct that includes the discriminator domain, the binding kinetics of tRNA^Ile^ to both the decoding and discriminator domains of *Mtb ileS* T-box riboswitch are revealed, significantly expanding our understanding of the tRNA binding mechanism that leads to genetic switching. This work illuminates how the interplay of structural elements in T-box riboswitches leads to unique conformational dynamics in different T-box riboswitches with different roles in biological regulation.

By using smFRET characterization with the same fluorophore pair at equivalent positions in the T-box and tRNA ligand as in our previous studies on the *B. subtilis glyQS* T-box (19), the current study allows comparison of tRNA binding mechanisms between transcription- and translation-regulating T-box riboswitches. The data support a similar two-step binding mechanism for the *Mtb ileS T-box* and *B. subtilis glyQS* T-box riboswitches, in which anticodon recognition by the decoding module occurs first, leading to the formation of a partially bound state, followed by binding of the tRNA NCCA end to the discriminator module to form the fully bound state. In both cases, the fully bound states are very stable, with a lifetime of at least 20 s, to secure sufficient time for transcriptional or translational outputs.

Besides the general similarity in the two major binding states, the data reveal unique features associated with the partially bound state of the *Mtb ileS* T-box riboswitch. In a previous study using a mutant of the *B. subtilis glyQS* T-box lacking the discriminator domain to specifically probe the partially bound state using tRNA^Gly^, it was found that the partially bound state in the *B. subtilis glyQS* T-box complex homogenously exhibited a short lifetime (∼ 3.6 s) before the tRNA dissociated (19). In contrast, the partially bound state in the *Mtb ileS* T-box complex has a 5-fold increase in the average lifetime (17.6 s) using a similar construct. The difference in the stability of the partially bound state in these two types of T-box riboswitches may be functionally linked to the temporal constraints of their regulations. In the case of *B. subtilis glyQS* T-box, an uncharged tRNA needs to fully bind before transcription and folding of the more thermostable terminator in order to lock the antiterminator configuration. The time needed for this decision is short. Therefore, a short lifetime of the partially bound state can avoid trapping of the *glyQS* T-box-tRNA complex by a charged tRNA into the terminator state, and potentially allows multiple sampling events of tRNA ligands. In the case of *Mtb ileS T-box*, translation regulation is not limited to co-transcriptional regulation, but could also potentially occur on fully transcribed mRNAs. In this context, multiple samplings of tRNA ligands are not limited to a short co-transcriptional time window, therefore a short lifetime of the partially bound state does not provide any functional benefits. On the other hand, a longer lifetime of the partially bound state may allow sufficient time for establishment of the NCCA-discriminator interactions downstream.

In addition, the data reveals conformational heterogeneity in the partially bound state in the *Mtb ileS* T-box riboswitch, which is absent in the *B. subtilis glyQS* T-box. When using a tRNA lacking the NCCA sequence, around 30% of the population forms a partially bound state with an extended lifetime and with rapid transitions into a state which resembles the fully bound state when using an uncharged intact tRNA. However, without being able to establish the NCCA-discriminator interactions, the lifetime of the 0.7 FRET state is two orders of magnitude shorter than the functional fully bound state. While the exact molecular architectures associated with this conformational heterogeneity remain unclear, the observation suggests that the *Mtb ileS T-box* riboswitch can dynamically sample the fully bound conformation to attempt interactions between the tRNA NCCA end and the discriminator module, and that the presence of the uncharged tRNA NCCA end can greatly stabilize this conformation. This conformation selection model of tRNA NCCA binding by the *Mtb ileS* T-box riboswitch is further supported by the smFRET measurement using the IntraFRET construct. Contrary to the case in the *B. subtilis glyQS* T-box, in which the intra-T-box conformational change involves an inward movement of the discriminator module relative to the decoding domain specifically accompanied by the establishment of NCCA-discriminator interactions (19), the *Mtb ileS* T-box displays a more relaxed state (zero FRET) and a more compact (0.3 state FRET) in the absence of tRNA ligand. Binding of tRNA^Ile^ and tRNA^Ile-ΔNCCA^ can both shift the equilibrium to the more compact conformation, with the former being more effective than the latter. Taken together, these findings support the notion that while binding of the tRNA NCCA to the *B. subtilis glyQS* T-box is likely to follow an induced-fit model, binding of tRNA NCCA to the *Mtb ileS* T-box is likely to follow a conformation selection model. This observation is crucial to understand the differences in biological function for the two types of T-box riboswitches. Transcription-regulating T-boxes need to make a rapid decision to allow or prevent transcription. This mechanistically favors a pre-formed, binding competent conformation of the riboswitch. Without such time limit, translation-regulating T-boxes are able to have a longer lifetime in the partially bound state, enabling the use of a conformation selection mechanism that rapidly samples the fully bound state, which becomes stabilized and consequently selected by NCCA-discriminator interactions.

Finally, our results highlight two critical sequence elements that facilitate tRNA binding and NCCA end discrimination. First, previous studies emphasized the central role of the Stem IIA/B-linker pseudoknot as a topological hinge that controls the conformations of Stems I and II (9). Consistent with this previous finding, the data from the Linker Mutant further illustrates that the A-minor interaction and the pseudoknot structure are critical for the stability of both the partially bound state and the fully bound state. The critical importance of these interactions helps explain the conservation of the ‘F-box’ sequence in T-boxes (nucleotides 77-81, CGUCA in Mtb ileS T-box) (29). Second, the present data elucidates the kinetic impact of the conserved RAG sequence on NCCA discrimination. Structural studies demonstrate that the RAG motif establishes specific interactions with the uncharged tRNA NCCA end. The results here demonstrate that mutating the RAG motif leads to the formation of a pseudo fully bound state, in which the tRNA NCCA end interacts with the discriminator domain with a distinct, likely non-functional configuration. This observation indicates that the RAG motif provides specificity for NCCA binding by constraining the conformational flexibility and reducing sampling of an aberrant NCCA docking state. While no equivalent measurement is available for a transcription-regulating T-box, given the structural similarity of the core discriminator domain and tRNA, it is possible to speculate that the conserved RAG motif may function in the same way in transcription regulating T-boxes as in translation regulating ones.

In summary, the current work together with previous kinetic characterization on the decoding domain (23,24) provides a more complete picture of tRNA ligand binding process by translational T-box riboswitches (**Figure 8**). All T-box riboswitches studied to date follow a two-step binding mechanism with anticodon recognition first, followed by establishment of the NCCA-discriminator interactions. In the case of translation-regulating T-boxes, the apo state samples an open and a closed conformation, the latter probably resembling the tRNA bound state. The decoding module of translational T-box riboswitches exhibits different conformations, and binding of the tRNA anticodon induces formation of a more compact and stable binding state (23,24), which likely corresponds to the partially bound state in this study. After anticodon recognition, tRNA in the partially bound state can transiently dock to the discriminator domain even in the absence of the tRNA NCCA-discriminator interactions, highlighting a higher conformational flexibility in translation-regulating T-box riboswitches compared to the transcription-regulating ones. This observation points to a fundamental functional difference between the two types of T-box riboswitches that may help explain how they have adapted to different functional constraints. Establishment of the tRNA NCCA-discriminator interaction finally locks the complex into a stable, fully bound state to allow translation initiation. In addition, the A81:G68:C77 A-minor interaction and the pseudoknot structure formed by Stem IIA/B and the linker region critically coordinate with both binding steps and enhance the stability of the partially bound and fully bound states. Finally, the conserved RAG sequence constrains the binding pocket of the tRNA NCCA end and provides specificity to NCCA recognition by limiting conformational flexibility. Overall, these findings shed light on the sophisticated and dynamic nature of T-box-tRNA interactions, highlighting the complex interplay between structural motifs, conformational flexibility, and ligand binding kinetics. By characterizing the nuanced structural adaptations that enable novel conformational dynamics, these results provide a more comprehensive framework to understand how structured RNA-based regulatory systems adapt to new regulatory niches.

**Figure 8.**
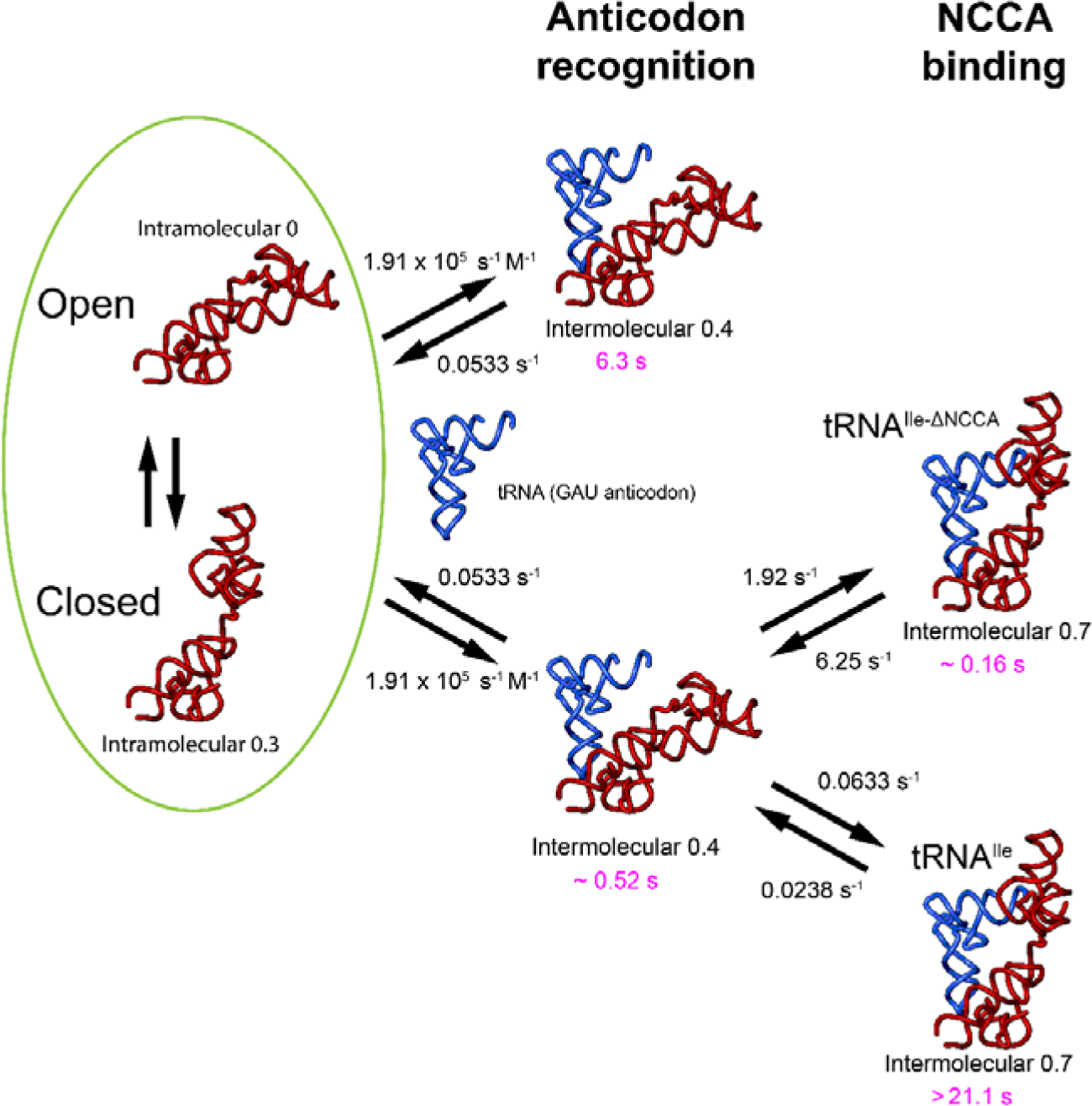
Proposed kinetic model for the binding process of the *Mtb-ileS* T-box to tRNA. In the absence of tRNA, the T-box fluctuates between a closed and an open conformation (left, green ellipse). In the presence of tRNA, the T-box binds tRNA through a two-step process where interactions with the anticodon are established first, forming a partially bound state (middle). The partially bound state exhibits conformational heterogeneity. One subpopulation (middle, top), with a 6.3 s lifetime, binds the tRNA anticodon but never transitions to the fully bound state before dissociating. The other subpopulation (middle, bottom) binds the tRNA anticodon and subsequently rapidly and continuously fluctuates between the partially and fully bound states, as in the absence of the NCCA end the interactions with the discriminator domain cannot be established. These fluctuations result in an unstable complex with a short lifetime resembling the fully bound state (right, top). The partially bound state with a short lifetime (middle, bottom), remains bound to tRNA for a relatively long time before full tRNA dissociation. In contrast, in the presence of an intact, uncharged NCCA end, the NCCA end of tRNA engages with the discriminator domain, leading to a long-lived fully bound state (right, bottom). Rate constants (black) are shown for the different binding states. Rates were extracted from Global HMM models except in the case of the rapid transitions between the short-lived partially bound state (middle, bottom) and fully bound state with tRNA^Ile-ΔNCCA^ (right, top). The GMM model used for this subpopulation precludes rate analysis, so rates were estimated as 1/τ. The rates for the first step were obtained from the tRNA^Ile-ΔNCCA^ mutant as this data reflects better the binding process when no interactions with the discriminator domain have been established. The rate constant from the unbound to the anticodon-bound state was calculated using a tRNA concentration of 100 nM. The lifetime of each of the states are shown in pink. The FRET efficiency value associated with each observed state is also shown.

## Methods

### Template DNA preparation

DNA templates, encompassing the sequence of interest placed downstream of a T7 promoter followed by a BsmI restriction site, were synthesized, and cloned into a pUC19 vector by GenScript (Jiangsu, China). For each plasmid, after transformation into *E. coli* DH5α cells, a minimum of 1 L of culture was grown overnight at 37°C shaking at 200 rpm in Lysogeny Broth (LB) supplemented with ampicillin (at a final concentration of 100 µg/ml). Cells were harvested by centrifugation at 5,470 x g for 30 minutes at 4°C. Each cell pellet from a 1 L culture was resuspended in 10 ml of GTE buffer (50 mM glucose, 25 mM tris-HCl pH 8.0, 10 mM EDTA, and 0.1 mg/ml RNase A), followed by addition of 20 ml of 0.2 M NaOH and 1% SDS to lyse the cells and left to stand on ice for five minutes. After neutralizing the lysate with 15 ml of 3 M potassium acetate (pH 5.2), the mixture was left on ice for ten minutes. In order to precipitate the DNA, the lysate supernatant was mixed with 0.6 volume of isopropanol followed by incubation at -20°C for 15 minutes. The solution was centrifuged for 30 minutes at 4°C at 4,000 x g and the supernatant was discarded. The DNA pellet was washed repeatedly with 80% ethanol and then resuspended in 2 ml of TE buffer (10 mM Tris pH 8.0, 0.1 mM EDTA). To eliminate RNA contamination, RNase A (4 mg/ml, final concentration 20 µg/ml) was added and incubated at 37°C for 20 minutes. Twice two equal volumes of phenol/chloroform (1:1) were used to extract the DNA, followed by addition of 10% (v/v) 3M sodium acetate pH 5.5 to precipitate the mixture. The precipitated DNA was rinsed twice with 80% ethanol before precipitation by centrifugation at 4,000 x g for 30 min at 4°C. After resuspension in 4 ml of TE buffer, 1.2 ml of 40% PEG 8,000 and 516 µl of 5 M NaCl were added, and the mixture was left to precipitate overnight at 4°C. The precipitated plasmid DNA was recovered by centrifugation at 4,000 x g for 30 min at 4°C, re-suspended in water, and ethanol precipitated again using 3 M sodium acetate pH 5.5 overnight at -20 °C. After three 80% ethanol washes, the pellet was once again resuspended in 1 ml of water. The concentration and purity of the purified plasmid DNA was assessed by measuring the absorbance ratio (A_260_/A_280_) using a Nanodrop spectrophotometer (Thermo Scientific) and aiming for a 1.9 ratio. The plasmids DNA were kept at 4 °C until needed. To confirm and validate the plasmid, the entire plasmid was sequenced (Primordium Laboratories).

### Fluorophore conjugation of DNA oligonucleotides

For intermolecular smFRET experiments, DNA oligonucleotides containing an amine modification at the 3’ end and a biotin modification at the 5’ end were synthesized by Integrated DNA Technologies (IDT, Coralville IA). For intramolecular smFRET experiments, an additional DNA oligonucleotide containing an amine modification at the 5’ end was synthesized by IDT. To attach either Cy5 or Cy3 to these oligos, 13.5 µl of the 100 µM DNA oligonucleotide was thoroughly mixed with 1.5 µl of 1 M NaHCO_3_ pH 8.6. Following this, a pre-prepared solution containing 25 µg of NHS-conjugated fluorophore (either Cy3 or Cy5 (Lumiprobe, Hunt Valley, MD)) dissolved in 0.5 µl DMSO was added to the DNA oligonucleotide mixture. The mixture was incubated at a constant 37°C temperature overnight to ensure proper conjugation. To precipitate the conjugated DNA oligonucleotide, 1.67 µl of 3 M sodium acetate pH 5.5 and 50 µl of pure ethanol were added to the mixture, followed by overnight incubation at −20°C. Post incubation, the mixture was centrifuged at 21,000 x g for 30 minutes. The resulting pellet, containing the conjugated DNA oligonucleotide, was then resuspended in 40 µl of water. To ensure the removal of any residual free dye and salt, the DNA solution was passed through a Micro Bio-Spin 6 column (Bio-Rad, Hercules, CA). The concentrations of oligo and conjugated fluorophore were measured using a Nanodrop spectrophotometer. The labeling efficiency for all labeled oligonucleotides were estimated to be at least 90%.

### T-box preparation

The RNA molecules were synthesized by *in vitro* transcription from a linearized DNA template. To linearize the template, plasmids purified as described above harboring the region of interest were digested by BsmI (NEB, Ipswich MA). The reaction mixture was prepared in a nuclease-free microcentrifuge tube, consisting of 100 µg of plasmid DNA, 500 units of BsmI, the appropriate volume of 10X rCutSmart™ Buffer, and nuclease-free water to bring the final concentration of enzyme to less than 5% (v/v). The tube was then incubated in a constant temperature water bath at 65°C, for 1 hour. Post incubation, the reaction was assessed for completion through gel electrophoresis. The digested product was then purified by two phenol-chloroform extractions, ethanol precipitated, and resuspended in nuclease-free water, and the concentration of the digested product was measured using a Nanodrop spectrophotometer.

For T-box sample preparation the samples were purified from the transcription reaction without any denaturing steps. Transcription reactions were set up in a total volume of 500 μl, containing 50 μg of linearized DNA template, 40 mM Tris-HCl pH 8.0, 10 mM MgCl_2_, 1 mM spermidine, 50 µg/ml BSA, 20 mM DTT, 5 mM of ATP, CTP, and UTP, 6 mM of GTP, 80 U of Recombinant RNase inhibitor (Promega, Fitchburg, WI), 8 U of inorganic pyrophosphatase (Sigma-Aldrich, St. Louis, MO), and 5 U of T7 RNA polymerase (purified in-house). A Cy3-labelled DNA oligonucleotide was added to a final concentration of 1 µM to the transcription reaction to promote co-transcriptional annealing to the 3’-extension of the T-box sequence. The mixture was gently mixed by pipetting and incubated at 37°C overnight. The next day, transcription reactions were briefly centrifuged to separate precipitated material before being loaded to a Superdex 200 10/200 GL column (Cytiva) preequilibrated with SEC buffer (10 mM Tris pH 7.4, 100 mM KCl and 20 mM MgCl_2_). The column was run with an isocratic flow at 0.5 ml/min, and fractions were collected between 11.5 and 14 ml, depending on the mutant (**Supplementary** Fig. 1). Collected fractions were concentrated to ∼40 µl by ultrafiltration using a Vivaspin 500 (Vivaproducts, Littleton, MA) centrifugal concentrator with a 3 kDa MWCO. The concentration of Cy3 in the concentrated fractions was measured using a Nanodrop spectrophotometer. One volume of 50% glycerol was subsequently added to the concentrated fractions before aliquoting, flash-freezing in liquid nitrogen, and storing at -80°C.

### tRNA preparation

For tRNA sample preparation, transcription reactions were set up in a total volume of 5 ml, containing 500 μg of linearized DNA template, 40 mM Tris-HCl pH 8.0, 10 mM MgCl₂, 1 mM spermidine, 50 µg/mL BSA, 20 mM DTT, 5 mM of ATP, CTP, and UTP, 6 mM of GTP, 800 U of Recombinant RNase inhibitor (Promega, Fitchburg, WI), 8 U of inorganic pyrophosphatase (Sigma-Aldrich, St. Louis, MO), and 5 U of T7 RNA polymerase (purified in-house). The mixture was gently mixed by pipetting and incubated at 37°C overnight. One volume of 50% glycerol was subsequently added to the transcription reaction before aliquoting, flash-freezing in liquid nitrogen, and storing at -80°C. Transcription samples were purified by urea-polyacrylamide gel electrophoresis (urea-PAGE) by cutting out the tRNA band. The excised gel slice was electroeluted using an EluTrap device (Whatman). After electroelution, tRNA was precipitated by the addition of 3 volumes of 100% ethanol and 1/10th volume of 3 M sodium acetate (pH 5.5), followed by incubation at -20°C for at least 1 hour. The tRNA was then pelleted by centrifugation at 12,000 x g for 15 minutes at 4°C, and the supernatant was discarded. The tRNA pellet was washed twice with 70% ethanol, centrifuged again at 12,000 x g for 5 minutes at 4°C. The tRNA pellet was dried using a SpeedVac vacuum concentrator (Savant) for 2–5 minutes and resuspended in an appropriate volume of nuclease-free water. The concentration of the purified RNA were assessed using a Nanodrop spectrophotometer. The purified tRNA was aliquoted and flash-frozen using liquid nitrogen and stored at -80°C until further use.

For 5’ end labelling of tRNA, the N-(3-Dimethylaminopropyl)-N′-ethylcarbodiimide hydrochloride (EDC) – N-hydroxysuccinimide (NHS) coupling method (30) was followed, wherein the 5’ monophosphate of the RNA is activated by EDC and imidazole for subsequent labelling. First the 5’ triphosphate of RNA was converted into a 5’ monophosphate by incubating 100 µg RNA with 100 units of RNA 5’ Pyrophosphohydrolase (NEB, Ipswich MA) at 37°C for 1 hour within a 100 µl reaction volume. Following this, the enzyme was removed by phenol chloroform extraction followed by passing the supernatant through a Micro Bio-Spin 6 column to exchange to a 10 mM HEPES pH 7.0, 150 mM NaCl, and 10 mM EDTA buffer. After the buffer exchange, the RNA solution was treated with 12.5 mg of EDC, 50 µl of ethylene diamine, and 200 µl of a 0.1 M imidazole pH 6.0 buffer. This mixture was then incubated for 3 hours at constant 37°C temperature. Post incubation, the RNA was ethanol precipitated and the pellet was resuspended in 0.1 M sodium carbonate pH 8.7 buffer. Any residual EDC was removed by passing the solution through a Bio-Spin 6 column. For the final labelling step, the RNA solution in 0.1 M sodium carbonate pH 8.7 buffer was incubated with Cy5 NHS (Lumiprobe, Hunt Valley, MD) dye for 45 minutes, using an approximate RNA to dye molar ratio of 1:200 followed by ethanol precipitation and RNA resuspension in water. To remove any residual free dye, the final RNA solution was passed through a Micro Bio-Spin 6 column. The labeling efficiency of tRNAs was consistently around 90%-100%.

Refolding tRNA for smFRET or BLI experiments was done by initially diluting it to the required final concentration in a solution containing 10 mM HEPES pH 7.5 and 50 mM NaCl. The sample was then heated to 95°C using a thermocycler, incubated for 3 minutes, and then transferred to ice to cool it rapidly for 3 minutes. After cooling, MgCl_2_ was added to a final 15 mM concentration. The tRNA solution was heated to 50°C in a thermocycler and incubated for 10 minutes at this temperature, followed by a 30 min incubation at 37 °C. Lastly, the sample was removed from the thermocycler and allowed to equilibrate to room temperature. For BLI binding assays, the folded tRNAs were dialyzed into 10 mM Tris pH 7.4, 100 mM KCl, and 20 mM MgCl_2_. For smFRET studies, the tRNA solution was added with one volume of 50% glycerol added, aliquoted and flash frozen in liquid nitrogen for long-term storage prior to use.

### Biolayer interferometry assays

Biolayer Interferometry (BLI) was used to determine the equilibrium dissociation constant (K_D_) of Mtb-ileS T-box and tRNA^ile^ using streptavidin biosensors (SAX) in an Octet K2 system (FortéBio Inc. San Jose, CA). The assays were performed on black, flat-bottom 96-well microplates (Greiner Bio-One 655209) in a total volume of 200 μl with orbital shaking at 1,000 rpm. The instrument was controlled with the Data Acquisition 11.1 (FortéBio Inc. San Jose, CA) software package. Before each experiment, biosensors were hydrated in SEC buffer (10 mM Tris pH 7.4, 100 mM KCl and 20 mM MgCl_2_). At the start of each experiment, a baseline was established using SEC buffer for 30 seconds. Subsequently the T-box at a 7.5 ng/µL concentration in SEC buffer was bound to the streptavidin sensor for 250 seconds, followed by washing for 30 seconds with the same buffer to eliminate non-specific binding and establish a pre-association baseline. Next, the purified and refolded tRNA was introduced into the biosensor and the association signal was measured for up to 500 seconds, until apparent equilibrium was reached at the highest concentration. Finally, the dissociation curves were obtained by moving the biosensors back into wells with SEC buffer (**Supplementary Fig.1b**). Each BLI experiment was done at several different tRNA concentrations (measured before each experiment) in the 1.6 to 55.3 μM range. For each set of experiments, a sensor that was not loaded with any T-box was used as a negative control and reference well for later data subtraction. New streptavidin biosensors were used for each experiment.

The raw data was preprocessed using the Octet Data Analysis Software version 11.1 (FortéBio Inc. San Jose, CA). Preprocessing included Y-axis alignment, reference subtraction, and interstep correction to account for noise while sensors are moved between solutions. Three binding models were considered to describe the binding kinetics: a one-step binding model, a parallel one-step binding model, and a sequential two-step binding model. Data was fit to the three considered models using either the Octet Data Analysis Software version 11.1 (FortéBio Inc. San Jose, CA) for the one-step and parallel one-step binding models, or a custom software for the two-step binding model. To prepare the data for our custom model fitting software, preprocessed data files were exported from the Octet Data Analysis software and converted to plain text (.txt). This data was then processed using Python scripts in Jupyter notebooks to fit the data to the sequential two-step binding model. Jupyter notebooks containing the code used to fit and plot this data are provided in the Source Data. A detailed description of the mathematical model used to fit this data is provided in the Supplementary Information and the Jupyter Notebooks.

### smFRET measurement

Quartz slides and glass coverslips for total internal fluorescence microscopy were prepared according to previously published protocols (31,32). The surface of the slides and coverslips were coated with a mixture of poly-ethylene glycol (PEG, Mw = 500,000) and biotin-PEG (Mw = 500,000). Microfluidic channels were constructed between the slides and coverslips as previously described (31,32). The channels were further passivated with a solution of 10 µM BSA to prevent non-specific binding, followed by incubation with a solution containing 10 µM BSA and 1 μM streptavidin. An aliquot of folded T-box riboswitch, carrying the labeled and biotinylated DNA oligo, was diluted to a concentration of 0.5-1 nM, and immobilized to the surface of the chamber via biotin-streptavidin interactions. tRNA was diluted to a concentration of 100 nM in an imaging buffer comprising of 50 mM HEPES pH 7.0, 100 mM KCl, 15 mM MgCl_2_, 5 mM protocatechuic acid (PCA) (Sigma), 1 U/mL protocatechuate-3,4-dioxygenase (PCD) (MP Biomedical), and 2 mM Trolox (Sigma), and was flowed into the microfluidic channel. For the intramolecular smFRET experiments, the concentration of the dual labeled T-box RNA was 1 nM and the unlabeled tRNA concentration was either 5 or 13 μM. An objective based total internal reflection fluorescence (TIRF) microscope based on a Nikon Ti-E with 100X NA 1.49 CFI HP TIRF objective (Nikon) was used to perform the smFRET measurements. A 561 nm laser (Coherent Obis at a power density of 4.07 × 10^5^ W/cm^2^) and a 647 nm laser (Cobolt MLD at a power density of 5.88 × 10^5^ W/cm^2^) were used for Cy3 and Cy5 excitation respectively. Emissions from both donor and acceptor were passed through an emission splitter (OptoSplit III, Cairn), and collected at different locations on an EMCCD (iXon Ultra 888, Andor). For intermolecular FRET measurement, 1500 frames of time-lapse images were taken with 100 ms exposure time and 561 nm laser excitation. For the intramolecular FRET measurements, 5 frames of time-lapse images with 100 ms exposure time and 647 nm laser excitation were taken before the FRET recording using the 561 nm laser excitation. The biological samples (i.e. the in vitro transcribed tRNAs and T-boxes) were generated at least twice, each considered as a biological replicate. For each biological replicate, at least three technical replicates were done. In each measurement, T-box and tRNA were diluted, smFRET images were recorded, and analysis were performed independently. Before each smFRET experiment, Tetraspeck beads (ThermoFisher) were used to align the emission channels.

### Lifetime analysis

NIS Elements software was used to pick individual spots and generate intensity trajectories from the time-lapse images. For intermolecular FRET experiments, maximum intensity projection of Cy5 emission signal was generated, and signals above an intensity threshold were selected to generate regions of interest (ROIs). For the intramolecular FRET measurements, maximum intensity projection of Cy5 emission signal from the 5 frames with Cy5 direct excitation was generated, and signals above an intensity threshold were selected to generate ROIs. ROIs were projected into the Cy3 and Cy5 channels in the time-lapse images to generate Cy3 and Cy5 intensity trajectories. Fluorescent intensity trajectories were corrected for baseline and bleed-through in MATLAB as previously described (33). FRET traces were generated by calculating I_Cy5_ / (I_Cy5_+I_Cy3_) at each time point from the intensity trajectories.

Idealization of FRET trajectories was achieved using the global modeling algorithm in tMAVEN (26) unless otherwise specified. Since global modeling is equivalent to concatenating the ensemble of trajectories into a single, long trajectory which is then analyzed with an HMM, this approach is often blind to transiently sampled states. In cases of transient sampling, there are often not enough datapoints corresponding to a state for the algorithm to separate it. Therefore, to model the short-lived FRET states of the WT *Mtb ileS* T-box bound to tRNA^Ile-ΔNCCA^-Cy5, traces showing transient and stable sampling of the 0.5 FRET state were analyzed independently using a composite approach. Traces that sampled the 0.5 FRET state transiently were analyzed by a variational Bayesian algorithm clustered by K-means (composite HMM + K-means) (26). This approach infers an HMM for each trace and then clusters the inferred FRET means for each HMM using K-means, allowing it to capture the short lived 0.5 FRET events of each trace. Modeling of the traces that stably sampled the 0.5 FRET state was achieved using the variational Bayesian Gaussian mixture modelling (vbGMM) algorithm in tMAVEN (26). As this approach assumes that all the data points are generated from a mixture of a finite number of Gaussian distributions with unknown parameters, it is able to identify the short-lived 0.7 FRET state, since it is consistently sampled throughout the dataset. The vbGMM approach, however, precludes rate analysis, so rates were estimated as the inverse of the lifetime for each state (1/τ). When a global modeling approach was used, transition rates were extracted from the resulting transition matrix. In all other cases, lifetimes were calculated using single exponential fitting to the dwell time data for each FRET state using tMAVEN (26), except in the cases where only two FRET states were identified. For the latter cases, the lifetime of the bound state was calculated as 1/τ, τ being the transition rates between the two states.

## Data availability

The data supporting the findings of this study are available from the corresponding authors upon reasonable request.

## Acknowledgements

We thank Jiacheng Zhang for help with the initial experiments, Arabela Grigorescu for help with the BLI experiments, as well as other members of the Fei and Mondragón laboratory for help and suggestions. We thank Dr. Ruben Gonzalez (Columbia University) for sharing tMAVEN. Research was supported by the NIH (R35-GM118108 to A.M. and NIH Director’s New Innovator Award 1DP2GM128185 to J.F.). We acknowledge the help from the Northwestern University Keck Biophysics Facility. Support from the R.H. Lurie Comprehensive Cancer Center of Northwestern University to the Keck Biophysics Facility is acknowledged.

## Funding

Research was supported by the NIH (R35-GM118108 to A.M. and 1DP2GM128185 to J.F.).

## Notes

### Competing Interest Statement

The authors have declared no competing interest.

