## Supplemental material for "Translational T-box riboswitches bind tRNA by modulating conformational flexibility"

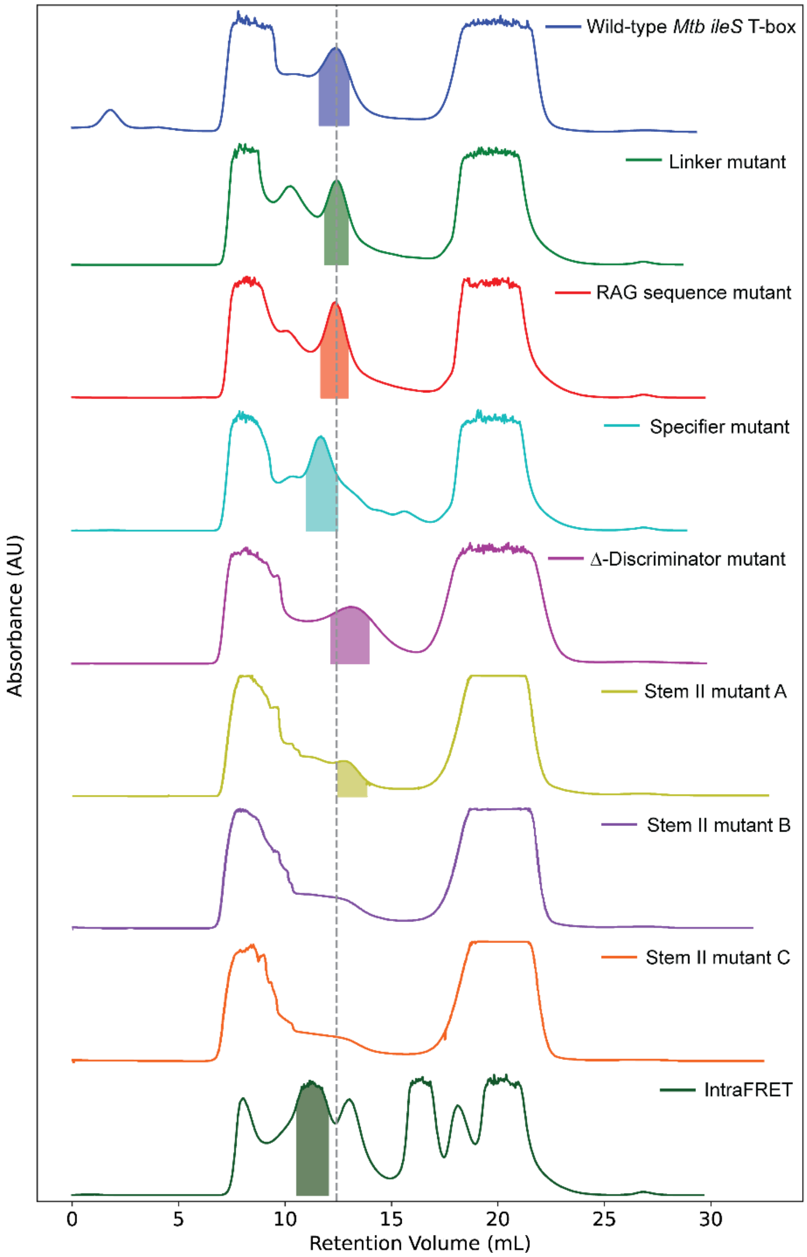

**Supplementary Figure S1. *Mtb-ileS* T-box riboswitch purification.** Size exclusion chromatography (SEC) elution profiles of the different constructs. The shaded regions correspond to the fractions that were selected for the FRET and BLI experiments. Slight variation in the elution of the T-box probably corresponds to changes in the hydrodynamic radius of the molecule. For Stem II Mutant B and C there was no clear peak detected, probably indicating that these mutants were misfolded. The T-box construct for intramolecular FRET measurement, with two dyes attached, elutes more rapidly reflecting the larger size of this molecule.

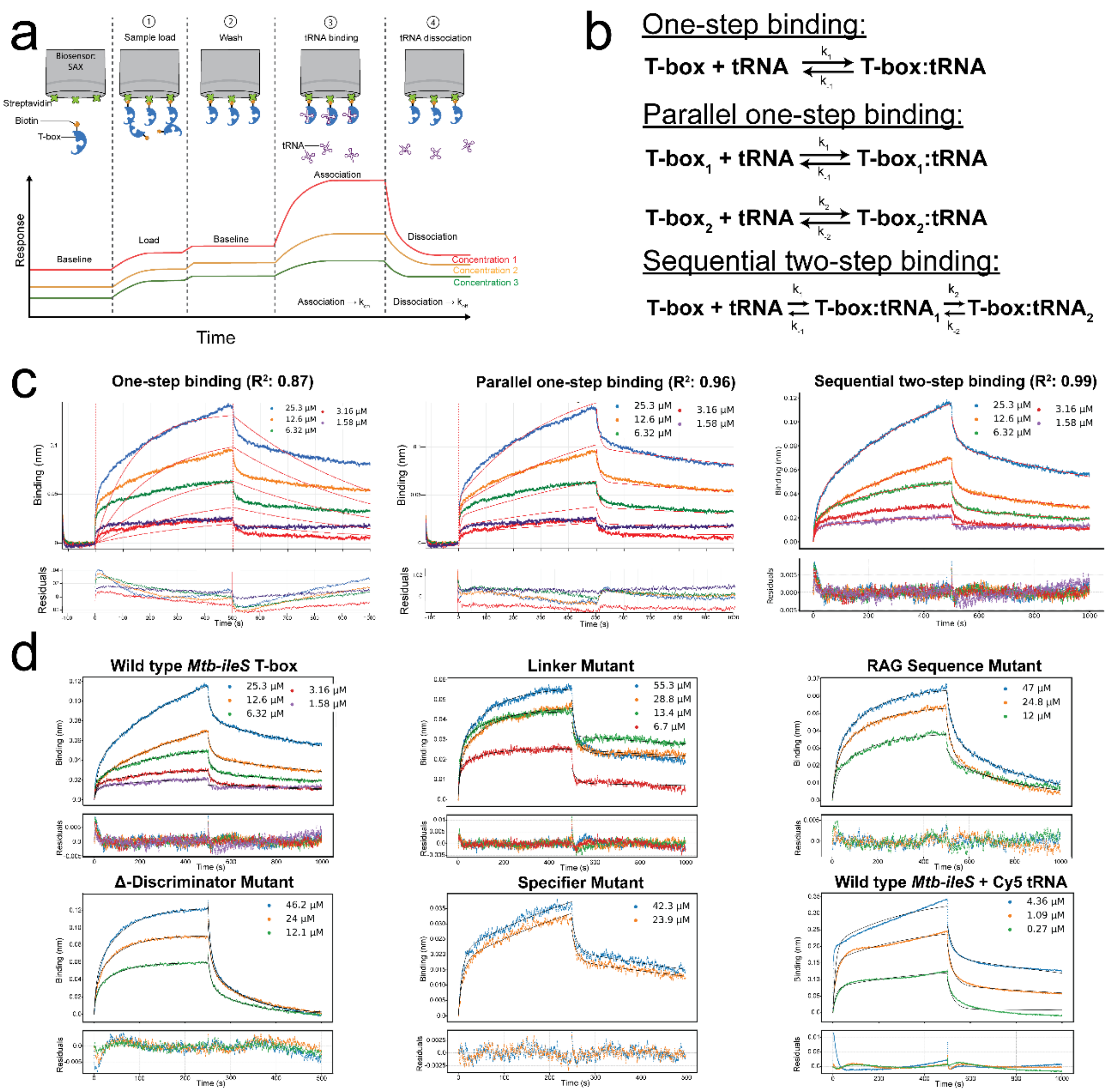

**Supplementary Figure S2. BLI analyses of the *Mtb-ileS* T-box riboswitch binding. a)** Schematic diagram showing the steps in the BLI experiment. In general, the experiment consists of four steps: loading of the labeled T-box, wash, binding of the tRNA (association step), and wash with buffer (dissociation step). During these steps, the response is measured against time. For analysis, only the last two steps, association and dissociation, are considered. **b)** Kinetic models considered to describe the BLI binding process: a one-step binding mechanism where the T-box and the tRNA bind in one single step, a parallel one-step binding scheme, where the two binding sites (anticodon:specifier and NCCA:T-box sequence) bind simultaneously but independently, and a sequential two-step binding scheme. **c)** BLI data on tRNA association and dissociation for the wild-type *Mtb-ileS* T-box was fit to the three kinetic models: one-step binding, parallel one-step binding, and sequential two-step binding. The sequential two step-binding model best fits the data ($R^2$ = 0.99) compared to the other models. The fits are shown by an orange dashed or solid line. The fit residuals are shown at the bottom. **d)** Fit of the BLI data for different constructs. In all cases, the sequential two-step binding model was used. The different tRNA concentrations used are shown by different colors and the concentrations are shown next to the plot. The fit is shown by a black solid line. The fit residuals are shown at the bottom. In some cases, like the specifier mutant, the binding was too weak to measure accurately more than a few concentrations. Descriptions of the binding model and the fitting procedure are found in the **Supplementary Methods** section. The fitting parameters are reported in **Supplementary Table I**.

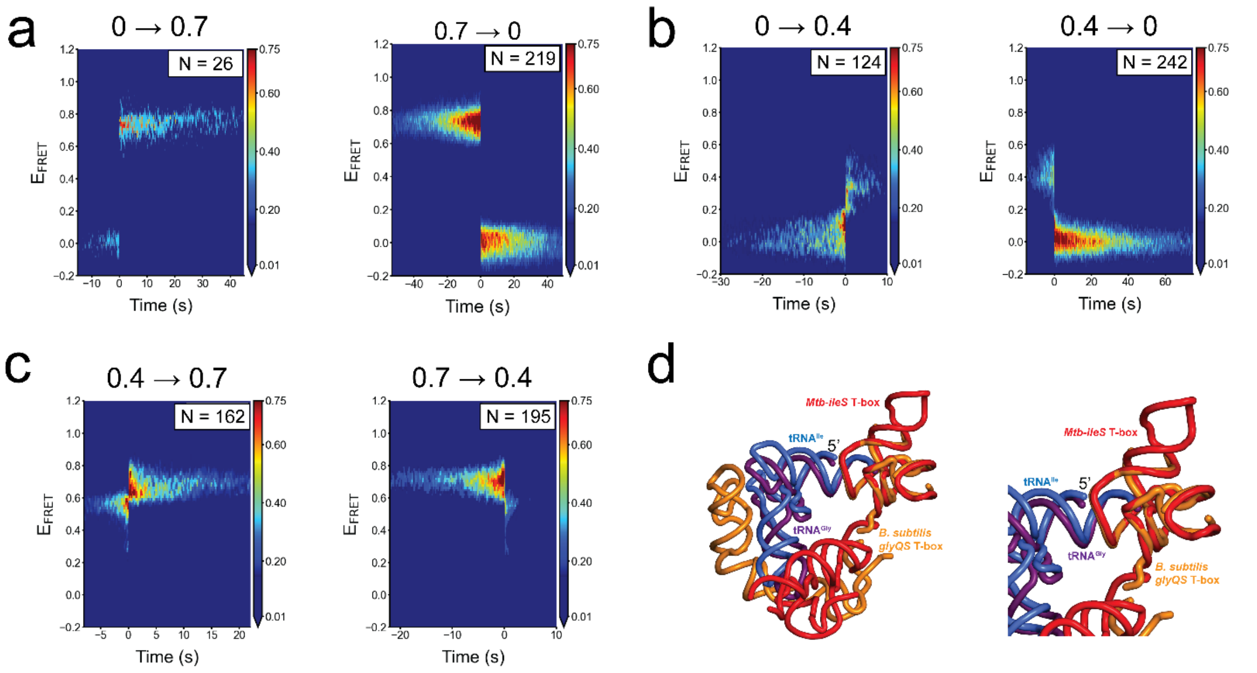

**Supplementary Figure S3. smFRET experiments of tRNA$^{Ile}$ binding to the wild-type *Mtb-ileS* T-box riboswitch show two distinct states. a)** Contour plots of the time-evolved FRET efficiency histogram displaying the transitions from the zero to 0.7 FRET state (left), and from 0.7 to zero FRET state (right). The data are synchronized with t = 0 corresponding to the transition point. N reports the number of transitions in each plot. The plots are contoured using normalized counts from 0.0 (blue) to 0.75 (red). **b)** Contour plots of the time-evolved FRET efficiency histogram displaying the transitions from the zero to 0.4 FRET state (left), and from 0.4 to zero FRET state (right). **c**) Contour plots of the time-evolved FRET efficiency histogram displaying the transitions from 0.4 to 0.7 FRET state (left), and from 0.7 to 0.4 FRET state (right). **d)** Superposition of the structure of *B. subtilis glyQS* T-box riboswitch/tRNA$^{Gly}$ complex (orange, purple) (1) on the *Mtb-ileS* T-box riboswitch/tRNA$^{ile}$ complex (red, blue) (2). The discriminator domains in the two structures were superposed to emphasize their similarity. The cartoon on the left shows the superposition of the two complexes, while the cartoon on the right shows a zoom of the discriminator domains. Overall, the superposition shows the high structural conservation of the discriminator domains and the way it interacts with the 3' end of the tRNA.

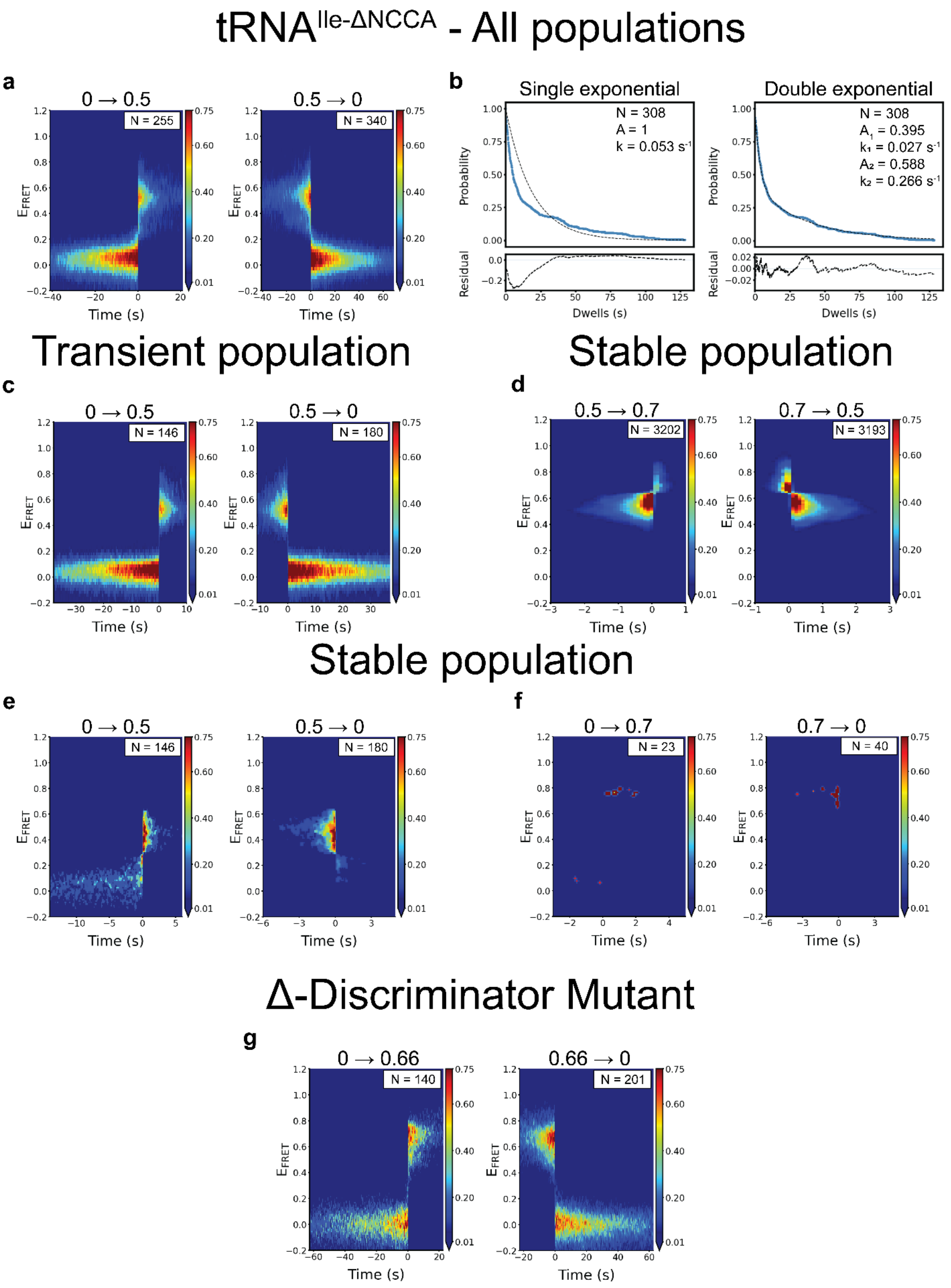

tRNAIle-ΔNCCA - All populations
a
0 → 0.5
N = 255
0.5 → 0
N = 340
EFRET
Time (s)
b
Single exponential
N = 308
A = 1
k = 0.053 s-1
Double exponential
N = 308
A1 = 0.395
k1 = 0.027 s-1
A2 = 0.588
k2 = 0.266 s-1
Probability
Residual
Dwells (s)
Transient population
c
0 → 0.5
N = 146
0.5 → 0
N = 180
Stable population
d
0.5 → 0.7
N = 3202
0.7 → 0.5
N = 3193
Stable population
e
0 → 0.5
N = 146
0.5 → 0
N = 180
f
0 → 0.7
N = 23
0.7 → 0
N = 40
Δ-Discriminator Mutant
g
0 → 0.66
N = 140
0.66 → 0
N = 201

**Supplementary Figure S4. smFRET experiments with mutants defective in discriminator binding help identify intermediate step. a)** Contour plots of the time-evolved FRET efficiency histogram displaying transitions from the zero to 0.5 FRET state (left), and from 0.5 to zero FRET state (right). The data are synchronized with t = 0 corresponding to the transition point. N reports the number of transitions in each plot. The plots are contoured using normalized counts from 0.0 (blue) to 0.75 (red). **b)** Dwell time survival plots for the 0.5 FRET state. The data were fitted using either a single ($Ae^{(-tk)}$, left) or double ($A_1e^{(-tk_1)} + A_2e^{(-tk_2)}$, right) exponential decay model. The double exponential decay model fits the data significantly better. The number of events (N) used for the analysis and the fitting parameters are shown. Data are shown in blue and the model in dotted black lines. The fit residuals are shown in the bottom plot. **c)** Contour plots of a time-evolved FRET efficiency histogram displaying transitions from the zero to 0.5 FRET state (left), and from 0.5 to zero FRET state (right) for the subpopulation of the *Mtb-ileS* T-box riboswitch/ $tRNA^{Ile\text{-}\Delta NCCA}$-Cy5 complex transiently sampling the bound state. **d)** Contour plots of a time-evolved FRET efficiency histogram displaying transitions from the 0.5 to 0.7 FRET state (left), and from 0.7 to 0.5 FRET state (right) for the subpopulation of the *Mtb-ileS* T-box riboswitch/ $tRNA^{Ile\text{-}\Delta NCCA}$-Cy5 complex stably sampling the bound state. **e)** Contour plots of a time-evolved FRET efficiency histogram displaying transitions from the zero to 0.5 FRET state (left), and from 0.5 to zero FRET state (right) for the subpopulation of the *Mtb-ileS* T-box riboswitch/ $tRNA^{Ile\text{-}\Delta NCCA}$-Cy5 complex stably sampling the bound state. **f)** Contour plots of a time-evolved FRET efficiency histogram displaying transitions from the zero to 0.7 FRET state (left), and from 0.7 to zero FRET state (right) for the subpopulation of the *Mtb-ileS* T-box riboswitch/ $tRNA^{Ile\text{-}\Delta NCCA}$-Cy5 complex stably sampling the bound state. **g)** Contour plots of a time-evolved FRET efficiency histogram of the *Mtb-ileS* T-box riboswitch Δ-Discriminator Mutant and $tRNA^{Ile}$ complex displaying transitions from zero to 0.66 FRET state (left) and 0.66 to zero FRET state (right).

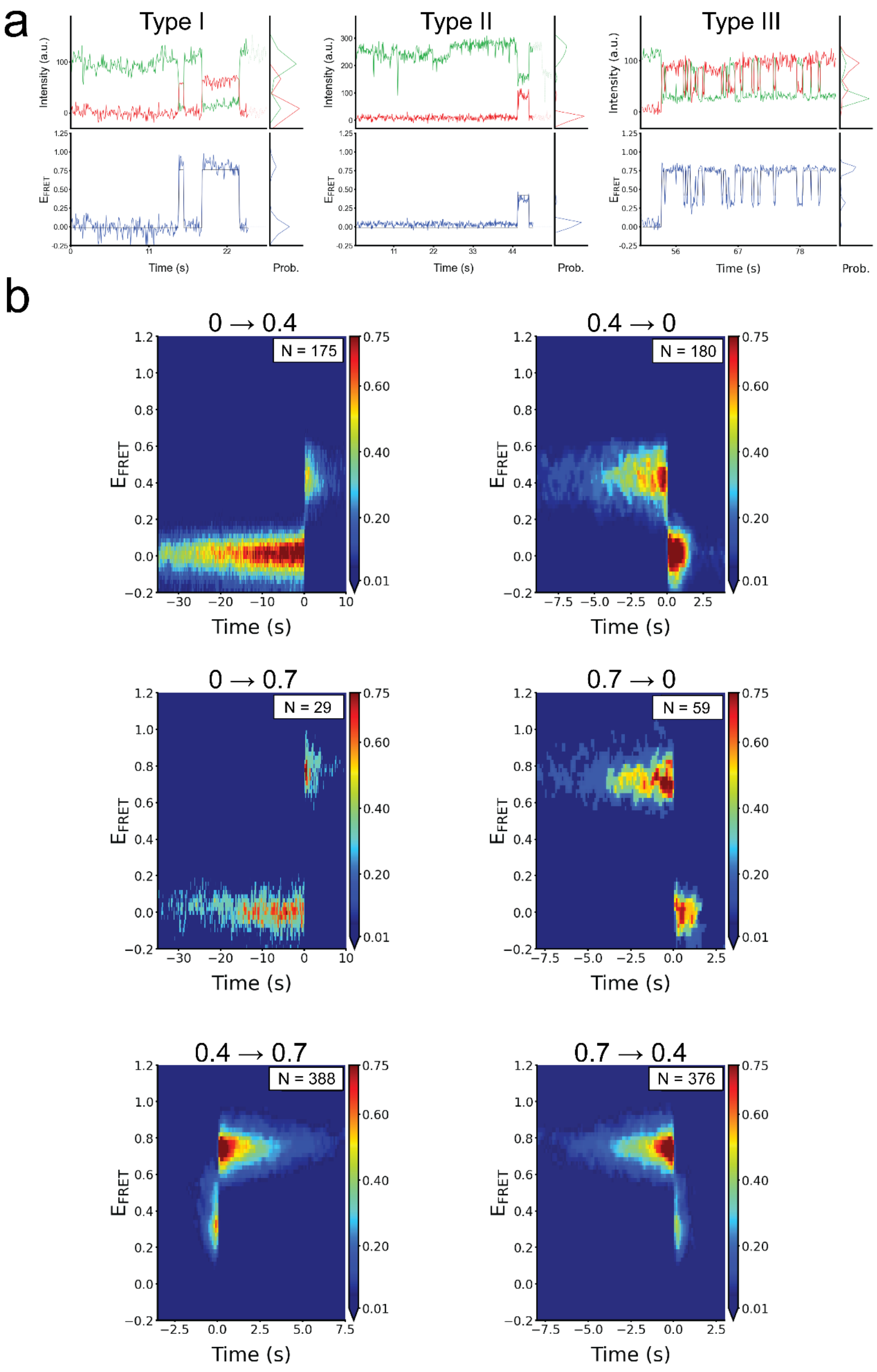

a
Type I
Type II
Type III
Intensity (a.u.)
$E_{FRET}$
Time (s)
Prob.
b
0 → 0.4
N = 175
0.4 → 0
N = 180
0 → 0.7
N = 29
0.7 → 0
N = 59
0.4 → 0.7
N = 388
0.7 → 0.4
N = 376
Time (s)

**Supplementary Figure S5. smFRET experiments with Linker Mutant show the importance of regions not directly contacting the NCAA sequence. a)** Representative Cy3 (green) and Cy5 (red) fluorescence intensity trajectories (top) and corresponding smFRET trajectories (bottom) for the Linker Mutant *Mtb-ileS* T-box riboswitch/tRNA$^{Ile}$ complex for the three types of traces identified. smFRET efficiency was calculated as ($I_{Cy5}$/($I_{Cy3}$-$I_{Cy5}$). **b)** Contour plots of a time-evolved FRET efficiency histogram of the Linker Mutant *Mtb-ileS* T-box riboswitch/ tRNA$^{Ile}$ complex displaying transitions between zero to 0.4 FRET state (top), between zero and 0.7 FRET state (middle), and between 0.4 and 0.7 FRET state (bottom). The data are synchronized with t = 0 corresponding to the corresponding transition point. N reports the number of transitions in each plot. The plots are contoured using normalized counts from 0.0 (blue) to 0.75 (red).

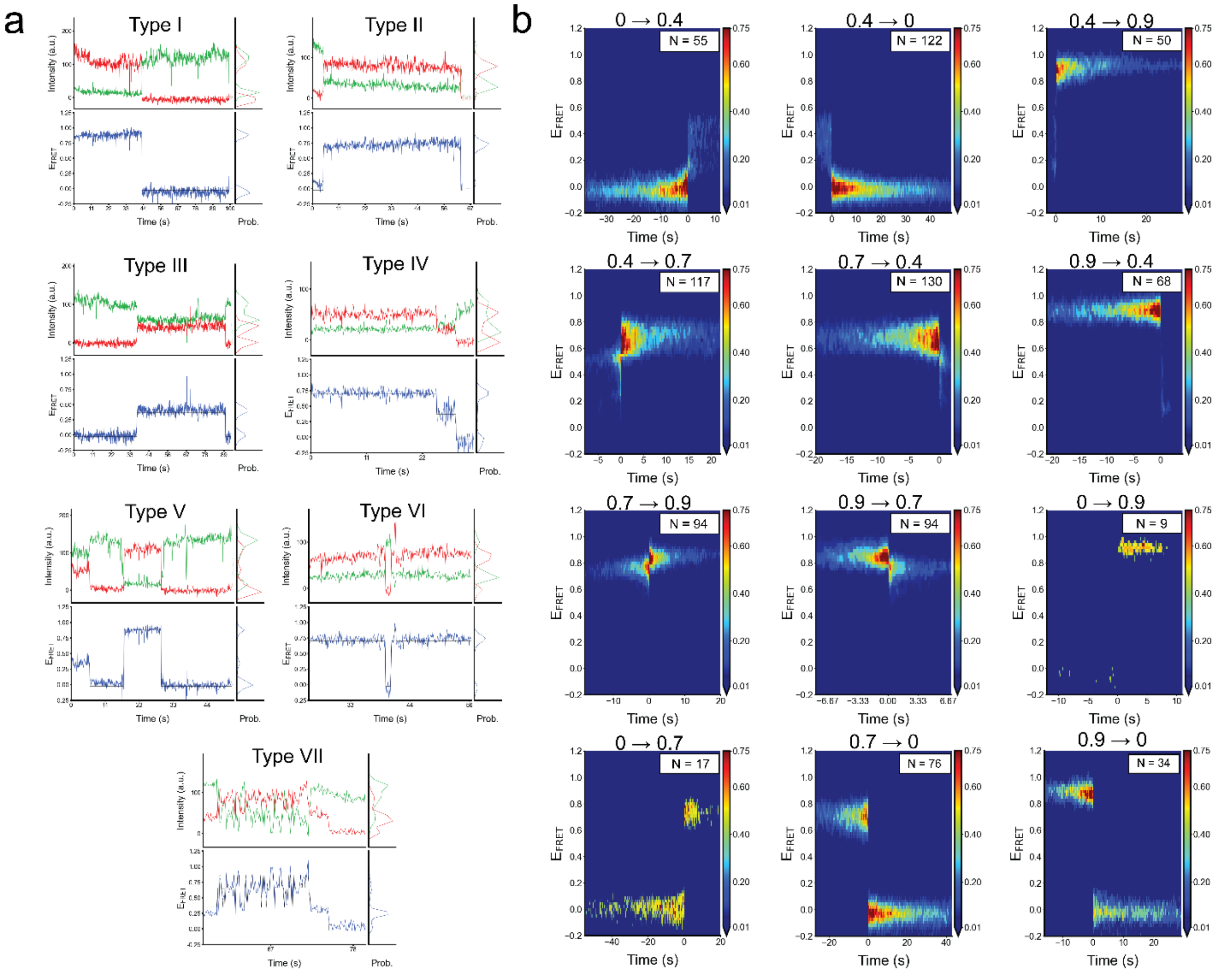

**Supplementary Figure S6. smFRET experiments with RAG Sequence mutant show the importance of contacts around the NCAA sequence. a)** Representative Cy3 (green) and Cy5 (red) fluorescence intensity trajectories (top) and corresponding smFRET trajectories (bottom) for the RAG Sequence mutant *Mtb-ileS* T-box riboswitch/tRNA$^{Ile}$ complex for all the types of traces identified. smFRET efficiency was calculated as ($I_{Cy5}/(I_{Cy3}$-$I_{Cy5}$). **b)** Contour plots of a time-evolved FRET efficiency histogram showing all transitions in the RAG Sequence mutant *Mtb-ileS* T-box riboswitch/ tRNA$^{Ile}$ complex. The data are synchronized with t = 0 corresponding to the corresponding transition point. N reports the number of transitions in each plot. The plots are contoured using normalized counts from 0.0 (blue) to 0.75 (red).

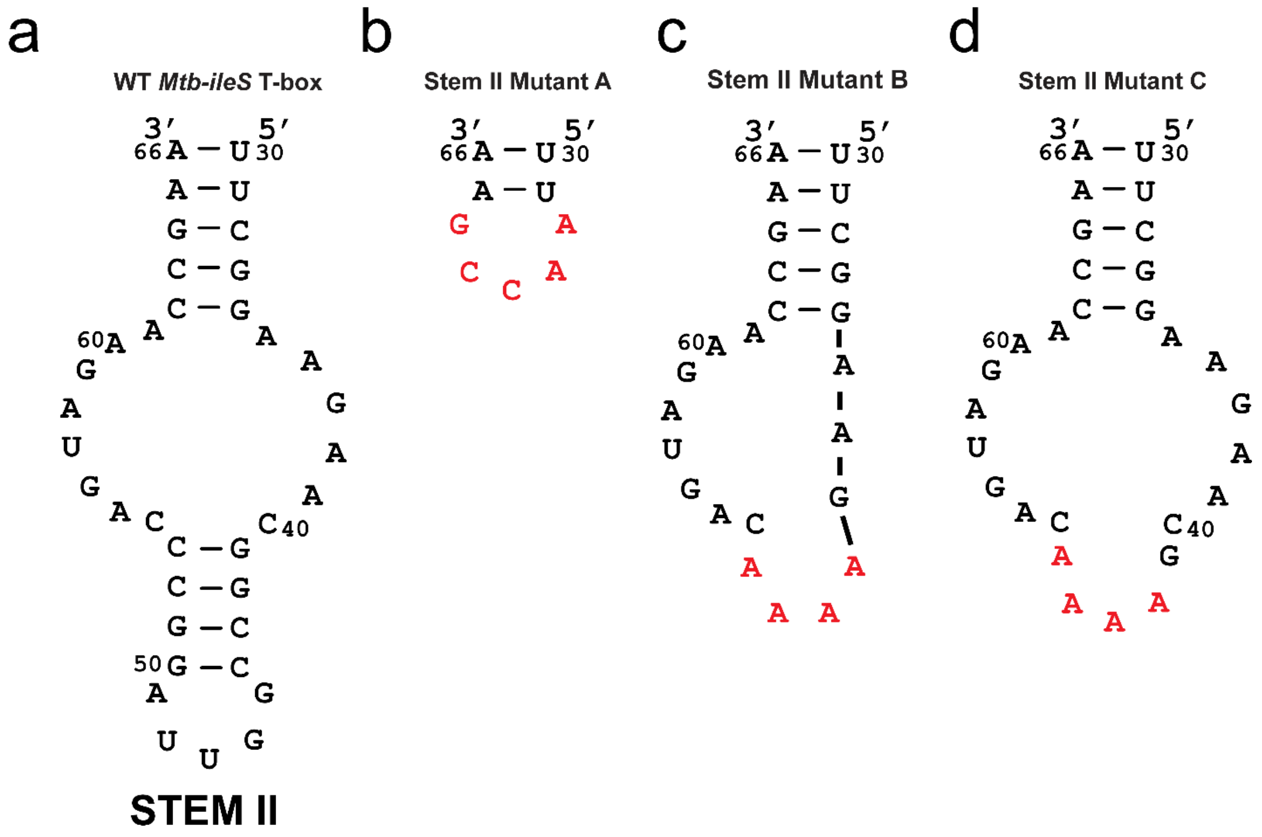

**Supplementary Figure S7. *Mtb-ileS* T-box riboswitch Stem II mutants analyzed. a)** Schematic diagram showing the sequence and paired regions in the wild-type Stem II in the *Mtb-ileS* T-box riboswitch. **b)** Schematic diagram of Stem II Mutant A where almost the entire Stem II (nucleotides 32-64) was deleted. **b)** Schematic diagram of Stem II Mutant B where one large region (nucleotides 38 -53), including the bottom paired region, was removed. **c)** Schematic diagram of Stem II Mutant C where the entire bottom paired region (nucleotides 42-53) was removed. In all diagrams new nucleotides introduced to form a loop are shown in red. Numbering according to the wild-type molecule.

| Biolayer interferometry (BLI) parameters fit to a two-step sequential model | | | | | | | |
|---|---|---|---|---|---|---|---|
| | $k_1$ ($M^{-1}\cdot s^{-1}$) | $k_{-1}$ ($s^{-1}$) | $k_2$ ($s^{-1}$) | $k_{-2}$ ($s^{-1}$) | $K_D$ (μM) | $R^2$ dissociation | $R^2$ association |
| **Wild type *Mtb* *Ile* T-box + Unlabeled tRNA** | 8781.53 | 0.0372 | 0.0072 | 0.0009 | 0.474 | 0.996 | 0.997 |
| **Wild type *Mtb* *Ile* T-box + Cy5 tRNA** | 5404.42 | 0.0352 | 0.0040 | 0.0008 | 1.03 | 0.985 | 0.980 |
| **Linker Mutant** | 4995.10 | 0.0350 | 0.0137 | 0.0004 | 0.182 | 0.975 | 0.991 |
| **RAG Mutant** | 10724.19 | 0.0537 | 0.0179 | 0.0075 | 1.48 | 0.984 | 0.993 |
| **Specifier Mutant** | 122.07 | 0.0882 | 0.0048 | 0.0014 | 160 | 0.917 | 0.973 |
| **Δ-Discriminator Mutant** | 10459.30 | 0.1252 | 0.0244 | 0.0150 | 4.55 | 0.993 | 0.997 |

**Supplementary Table I. Parameters fitted to the BLI data.** The table shows the fitting parameters and $R^2$ residual for tRNA association and dissociation for each construct. The sequential two-step binding model was used to fit the BLI data. The fit was done as described in the **Supplementary Methods** section. A diagram showing the fit to the data for the different constructs is shown in **Supplementary Figure S2.**

| Sample | Mean | Variance | Fraction |
|---|---|---|---|
| **Wild-type *Mtb-ileS* T-box + tRNA$^{Ile}$** | | | |
| **State 1** | -0.0081 | 0.0095 | 0.5901 |
| **State 2** | 0.4212 | 0.0165 | 0.0990 |
| **State 3** | 0.7357 | 0.0087 | 0.3110 |
| **Wild-type *Mtb-ileS* T-box + tRNA$^{Ile\text{-}\Delta NCCA}$** | | | |
| **Stable** | | | |
| **State 1** | 0.0368 | 0.0104 | 0.2426 |
| **State 2** | 0.5235 | 0.0067 | 0.4941 |
| **State 3** | 0.6951 | 0.0120 | 0.2634 |
| **Transient** | | | |
| **State 1** | 0.0318 | 0.0060 | 0.8941 |
| **State 2** | 0.5332 | 0.0144 | 0.1059 |
| **General** | | | |
| **State 1** | 0.0284 | 0.0107 | 0.7010 |
| **State 2** | 0.5289 | 0.0260 | 0.2990 |
| ***Mtb-ileS* T-box Δ-Discriminator Mutant + tRNA$^{Ile}$** | | | |
| **State 1** | -0.0147 | 0.0150 | 0.7543 |
| **State 2** | 0.6674 | 0.0161 | 0.2457 |
| ***Mtb-ileS* T-box Linker mutant + tRNA$^{Ile}$** | | | |
| **State 1** | -0.0179 | 0.0115 | 0.7107 |
| **State 2** | 0.4093 | 0.0160 | 0.1181 |
| **State 3** | 0.7572 | 0.0139 | 0.1712 |
| ***Mtb-ileS* T-box RAG mutant + tRNA$^{Ile}$** | | | |
| **State 1** | -0.0291 | 0.0053 | 0.5667 |
| **State 2** | 0.3609 | 0.0235 | 0.1075 |
| **State 3** | 0.6950 | 0.0068 | 0.1882 |
| **State 4** | 0.8799 | 0.0045 | 0.1377 |

**Supplementary Table II. Parameters of the consensus HMM model fitted to the FRET histogram for the different constructs.**

| Sample | Trace type | Number | Fraction |
|---|---|---|---|
| **Wild-type *Mtb-ileS* T-box + tRNA$^{ile}$** | | | |
| | I | 269 | 0.7730 |
| | II | 62 | 0.1782 |
| | III | 17 | 0.0488 |
| **Total** | | 348 | 1.0 |
| **Type I subtypes** | I-1 | 247 | 0.9182 |
| | I-2 | 22 | 0.0818 |
| **Total** | | 269 | 1.0 |
| **Wild-type *Mtb-ileS* T-box + tRNA$^{ile\text{-}\Delta NCCA}$** | | | |
| | Stable | 72 | 0.2727 |
| | Transient | 192 | 0.7273 |
| **Total** | | 264 | 1.0 |
| ***Mtb-ileS* T-box Δ-Discriminator Mutant + tRNA$^{Ile}$** | | | |
| | All | 161 | 1.0 |
| **Total** | | 161 | 1.0 |
| ***Mtb-ileS* T-box Linker mutant + tRNA$^{Ile}$** | | | |
| | I | 40 | 0.2030 |
| | II | 91 | 0.4620 |
| | III | 66 | 0.3350 |
| **Total** | | 197 | 1.0 |
| **Type I subtypes** | I-1 | 8 | 0.2 |
| | I-2 | 32 | 0.8 |
| **Total** | | 40 | 1.0 |
| ***Mtb-ileS* T-box RAG mutant + tRNA$^{Ile}$** | | | |
| | I | 72 | 0.3654 |
| | II | 35 | 0.1776 |
| | III | 17 | 0.0862 |
| | IV | 55 | 0.2791 |
| | V | 4 | 0.0203 |
| | VI | 8 | 0.0406 |
| | VII | 6 | 0.0304 |
| **Total** | | 197 | 1.0 |

**Supplementary Table III. Categorization of smFRET trace types.** The table shows the trace number and fraction of each trace type for different constructs. Refer to the text for a detailed description.

| Sample | Transition | Number of events | Fraction of events |
|---|---|---|---|
| **Wild-type *Mtb-ileS* T-box + tRNA$^{Ile}$** | | | |
| | 0 → 0.7 | 26 | 0.0269 |
| | 0 → 0.4 | 124 | 0.1281 |
| | 0.4 → 0 | 242 | 0.2500 |
| | 0.4 → 0.7 | 162 | 0.1674 |
| | 0.7 → 0 | 219 | 0.2262 |
| | 0.7 → 0.4 | 195 | 0.2014 |
| **Total** | | 968 | 1.0 |
| **Wild-type *Mtb-ileS* T-box + tRNA$^{Ile\text{-}\Delta NCCA}$** | | | |
| **Stable** | 0 → 0.5 | 230 | 0.0331 |
| | 0 → 0.7 | 23 | 0.0033 |
| | 0.5 → 0 | 250 | 0.0360 |
| | 0.5 → 0.7 | 3202 | 0.4615 |
| | 0.7 → 0 | 40 | 0.0058 |
| | 0.7 → 0.5 | 3193 | 0.4602 |
| **Total** | | 6938 | 1.0 |
| **Transient** | 0 → 0.5 | 146 | 0.4478 |
| | 0.5 → 0 | 180 | 0.5522 |
| **Total** | | 326 | 1.0 |
| **All populations** | 0 → 0.5 | 255 | 0.4286 |
| | 0.5 → 0 | 340 | 0.5714 |
| **Total** | | 595 | 1.0 |
| ***Mtb-ileS* T-box Δ-Discriminator Mutant + tRNA$^{Ile}$** | | | |
| | 0 →0.66 | 140 | 0.4106 |
| | 0.66 → 0 | 201 | 0.5894 |
| **Total** | | 341 | |
| ***Mtb-ileS* T-box Linker mutant + tRNA$^{Ile}$** | | | |
| | 0 → 0.7 | 29 | 0.0240 |
| | 0 → 0.4 | 175 | 0.1450 |
| | 0.4 → 0 | 180 | 0.1491 |
| | 0.4 → 0.7 | 388 | 0.3215 |
| | 0.7 → 0 | 59 | 0.0489 |
| | 0.7 → 0.4 | 376 | 0.3115 |
| **Total** | | 1207 | 1.0 |
| ***Mtb-ileS* T-box RAG mutant + tRNA$^{Ile}$** | | | |
| | 0 → 0.4 | 55 | 0.0635 |
| | 0 → 0.7 | 17 | 0.0196 |
| | 0 → 0.9 | 9 | 0.0104 |
| | 0.4 → 0 | 122 | 0.1409 |
| | 0.4 → 0.7 | 117 | 0.1351 |
| | 0.4 → 0.9 | 50 | 0.0577 |
| | 0.7 → 0 | 76 | 0.0878 |
| | 0.7 → 0.4 | 130 | 0.1501 |
| | 0.7 → 0.9 | 94 | 0.1086 |
| | 0.9 → 0 | 34 | 0.0393 |
| | 0.9 → 0.4 | 68 | 0.0785 |
| | 0.9 → 0.7 | 94 | 0.1086 |
| **Total** | | 866 | 1.0 |

**Supplementary Table IV. Number of transition events identified in different smFRET experiments.** The table shows the number of transitions identified and corresponding fraction in each experiment. Refer to the text for a detailed description.

| Sample | Transition rate matrix |
|---|---|

**Wild-type *Mtb-ileS* T-box + tRNA$^{Ile}$**

Final State (columns) / Initial State (rows)

| | 0.0 | 0.4 | 0.7 |
|---|---|---|---|
| **0.0** | - | 0.00741 | 0.00195 |
| **0.4** | 0.08036 | - | 0.06327 |
| **0.7** | 0.02367 | 0.02380 | - |

**Wild-type *Mtb-ileS* T-box + tRNA$^{Ile\text{-}\Delta NCCA}$**

Final State (columns) / Initial State (rows)

| | 0.0 | 0.5 |
|---|---|---|
| **0.0** | - | 0.01905 |
| **0.5** | 0.05326 | - |

***Mtb-ileS* T-box Δ-Discriminator Mutant + tRNA$^{Ile}$**

Final State (columns) / Initial State (rows)

| | 0.0 | 0.66 |
|---|---|---|
| **0.0** | - | 0.01340 |
| **0.66** | 0.05684 | - |

***Mtb-ileS* T-box Linker mutant + tRNA$^{Ile}$**

Final State (columns) / Initial State (rows)

| | 0.0 | 0.4 | 0.7 |
|---|---|---|---|
| **0.0** | - | 0.03326 | 0.00294 |
| **0.4** | 0.20843 | - | 0.52650 |
| **0.7** | 0.02478 | 0.36332 | - |

***Mtb-ileS* T-box RAG mutant + tRNA$^{Ile}$**

Final State (columns) / Initial State (rows)

| | 0.0 | 0.4 | 0.7 | 0.9 |
|---|---|---|---|---|
| **0.0** | - | 0.01272 | 0.00214 | 0.00127 |
| **0.4** | 0.12160 | - | 0.10789 | 0.03942 |
| **0.7** | 0.02947 | 0.06497 | - | 0.06125 |
| **0.9** | 0.02065 | 0.04484 | 0.07729 | - |

**Supplementary Table V. Transition rate matrices calculated by the consensus HMM modeling for different constructs.** The units of all the rates are $s^{-1}$. Refer to the text for a description of the calculations.

| Sample | FRET state | Lifetime (s) | Error (±) |
|---|---|---|---|
| **Wild-type *Mtb-ileS* T-box + tRNA$^{Ile}$** | | | |
| | 0.4 | 6.9 | 0.14 |
| | 0.7 | 21.1 | 0.09 |
| **Wild-type *Mtb-ileS* T-box + tRNA$^{Ile\text{-}\Delta NCCA}$** | | | |
| **All populations** | 0.5 | 18.8 | 0.34 |
| **Stable** | 0.5 | 0.52 | 0.01 |
| | 0.7 | 0.16 | 0.004 |
| **Transient** | 0.5 | 6.3 | 0.04 |
| ***Mtb-ileS* T-box Δ-Discriminator Mutant + tRNA$^{Ile}$** | | | |
| | 0.5 | 17.6 | 0.12 |
| ***Mtb-ileS* T-box Linker mutant + tRNA$^{Ile}$** | | | |
| | 0.4 | 0.63 | 0.01 |
| | 0.7 | 2.6 | 0.01 |
| ***Mtb-ileS* T-box RAG mutant + tRNA$^{Ile}$** | | | |
| | 0.4 | 7.5 | 0.22 |
| | 0.7 | 9.7 | 0.20 |
| | 0.9 | 8.8 | 0.10 |

**Supplementary Table VI. Lifetimes for the states observed for the different constructs.**

| Part | Insert sequence |
|---|---|
| **WT *Mtb IleS* T-box** | TCTAGATTTTAATACGACTCACTATAGGCATCGATCCGGCGATCACCGGGGAGCCTTC<br>GGAAGAACGGCCGGTTAGGCCCAGTAGAACCGAACGGGTTGGCCCGTCACAGCCTCA<br>AGTCGAGCGGCCGCGCATCGGCGTGGCAAGCGGGGTGGTACCGCGGCGTTCGCGCA<br>CCGGCGTGGCGTCGTCCCCGAAATGTTTGCGGCTGCAGCATTCTTTGCATGC |
| **Linker Mutant** | TCTAGATTTTAATACGACTCACTATAGGCATCGATCCGGCGATCACCGGGGAGCCTTC<br>GGAAGAACGGCCGGTTAGGCCCAGTAGAACCGAACGGGTTGGCCCGTCGCACGGCC<br>AAGTCGAGCGGCCGCGCATCGGCGTGGCAAGCGGGGTGGTACCGCGGCGTTCGCGC<br>ACCGGCGTGGCGTCGTCCCCGAAATGTTTGCGGCTGCAGCATTCTTTGCATGC |
| **RAG Sequence Mutant** | TCTAGATTTTAATACGACTCACTATAGGCATCGATCCGGCGATCACCGGGGAGCCTTC<br>GGAAGAACGGCCGGTTAGGCCCAGTAGAACCGAACGGGTTGGCCCGTCACAGCCTCA<br>AGTCCGCCGGCCGCGCATCGGCGTGGCAAGCGGGGTGGTACCGCGGCGTTCGCGCA<br>CCGGCGTGGCGTCGTCCCCGAAATGTTTGCGGCTGCAGCATTCTTTGCATGC |
| **Specifier Mutant** | TCTAGATTTTAATACGACTCACTATAGGCATCGATCCGGCGTGGACCGGGGAGCCTTC<br>GGAAGAACGGCCGGTTAGGCCCAGTAGAACCGAACGGGTTGGCCCGTCACAGCCTCA<br>AGTCGAGCGGCCGCGCATCGGCGTGGCAAGCGGGGTGGTACCGCGGCGTTCGCGCA<br>CCGGCGTGGCGTCGTCCCCGAAATGTTTGCGGCTGCAGCATTCTTTGCATGC |
| **Δ-Discriminator Mutant** | TCTAGATTTTAATACGACTCACTATAGGCATCGATCCGGCGATCACCGGGGAGCCTTC<br>GGAAGAACGGCCGGTTAGGCCCAGTAGAACCGAACGGGTTGGCCCGTCACAGCCTCA<br>AGTCAATGTTTGCGGCTGCAGCATTCTTTGCATGC |
| **Stem II Mutant A** | TCTAGATTTTAATACGACTCACTATAGGCATCGATCCGGCGATCACCGGGGAGCCTTAA<br>CCGAACGGGTTGGCCCGTCACAGCCTCAAGTCGAGCGGCCGCGCATCGGCGTGGCA<br>AGCGGGGTGGTACCGCGGCGTTCGCGCACCGGCGTGGCGTCGTCCCCGAAATGTTT<br>GCGGCTGCAGCATTCTTTGCATGC |
| **Stem II Mutant B** | TCTAGATTTTAATACGACTCACTATAGGCATCGATCCGGCGATCACCGGGGAGCCTTC<br>GGAAGAAAAGTAGAACCGAACGGGTTGGCCCGTCACAGCCTCAAGTCGAGCGGCCGC<br>GCATCGGCGTGGCAAGCGGGGTGGTACCGCGGCGTTCGCGCACCGGCGTGGCGTCG<br>TCCCCGAAATGTTTGCGGCTGCAGCATTCTTTGCATGC |
| **Stem II Mutant C** | TCTAGATTTTAATACGACTCACTATAGGCATCGATCCGGCGATCACCGGGGAGCCTTC<br>GGAAGAACGAAAACAGTAGAACCGAACGGGTTGGCCCGTCACAGCCTCAAGTCGAGC<br>GGCCGCGCATCGGCGTGGCAAGCGGGGTGGTACCGCGGCGTTCGCGCACCGGCGT<br>GGCGTCGTCCCCGAAATGTTTGCGGCTGCAGCATTCTTTGCATGC |
| **IntraFRET** | GAATTCTTTTAATACGACTCACTATAGGGGAAATTAGGGGGGGCATCGATCCGGCGA<br>TCACCGGGGAGCCTTCGGAAGAACGGCCGGTTAGGCCCAGTAGAACCGAACGGGTTG<br>GCCCGTCACAGCCTCAAGTCGAGCGGCCGCGCATCGGCGTGGCAAGCGGGGTGGTA<br>CCGCGGCGTTCGCGCACCGGCGTGGCGTCGTCCCCGAAATGTTTGCGGCTGCTGAGA<br>CCTCTCTCTCTCGGATCC |
| **$tRNA^{Ile}$ WT** | TCTAGATTTTAATACGACTCACTATAGGGCCTATAGCTCAGGCGGTTAGAGCGCTTCGC<br>TGATAACGAAGAGGTCGGAGGTTCAAGTCCTCCTAGGCCCACCAAGCATTCTTTGCAT<br>GC |
| **$tRNA^{Ile}$ ΔNCCA** | TCTAGATTTTAATACGACTCACTATAGGGCCTATAGCTCAGGCGGTTAGAGCGCTTCGC<br>TGATAACGAAGAGGTCGGAGGTTCAAGTCCTCCTAGGCCCAGCATTCTTTGCATGC |
| **$tRNA^{Trp}$** | TCTAGATTTTAATACGACTCACTATAGGGGGTATAGTTTAATGGTAAAACGAAGGTCTC<br>CAAAACCTTTGATGTGGGTTCGATTCCTACTACCCCCACCAAGCATTCTTTGCATGC |
| **3’ Extension Oligo** | 5’-/Biosg/GCAGCCGCAAACATT/3AmMC6T/-3’ |

| | |
|---|---|
| **5’ Extension Oligo (for intramolecular experiments)** | 5’-/AmMC6/CCCCCTAATTTCCCCC-3’ |

**Supplementary Table VII. Sequences used for inserting constructs into cloning vector and for anchoring oligonucleotides.**

| Part | 5' Cloning site | 3' Cloning site | Linearization site | T7 Promoter sequence | Overhang for DNA probe annealing (3') | Overhang for DNA probe annealing (5') |
|---|---|---|---|---|---|---|
| **WT Mtb IleS T-box** | XbaI (TCTAGA) | SphI (GCATGC) | BsmI (GCATTC) | TAATACG ACTCACT ATA | AATGTTTGC GGCTGC | - |
| **Linker Mutant** | XbaI (TCTAGA) | SphI (GCATGC) | BsmI (GCATTC) | TAATACG ACTCACT ATA | AATGTTTGC GGCTGC | - |
| **RAG Sequence Mutant** | XbaI (TCTAGA) | SphI (GCATGC) | BsmI (GCATTC) | TAATACG ACTCACT ATA | AATGTTTGC GGCTGC | - |
| **Specifier Mutant** | XbaI (TCTAGA) | SphI (GCATGC) | BsmI (GCATTC) | TAATACG ACTCACT ATA | AATGTTTGC GGCTGC | - |
| **Δ-Discriminator Mutant** | XbaI (TCTAGA) | SphI (GCATGC) | BsmI (GCATTC) | TAATACG ACTCACT ATA | AATGTTTGC GGCTGC | - |
| **Stem II Mutant A** | XbaI (TCTAGA) | SphI (GCATGC) | BsmI (GCATTC) | TAATACG ACTCACT ATA | AATGTTTGC GGCTGC | - |
| **Stem II Mutant B** | XbaI (TCTAGA) | SphI (GCATGC) | BsmI (GCATTC) | TAATACG ACTCACT ATA | AATGTTTGC GGCTGC | - |
| **Stem II Mutant C** | XbaI (TCTAGA) | SphI (GCATGC) | BsmI (GCATTC) | TAATACG ACTCACT ATA | AATGTTTGC GGCTGC | - |
| **IntraFRET** | EcoRI (GAATTC) | BamHI (GGATCC) | BsaI (GAGACC) | TAATACG ACTCACT ATA | AATGTTTGC GGCTGC | GGGGGAAA TTAGGGGG |
| **tRNA$^{Ile}$ WT** | XbaI (TCTAGA) | SphI (GCATGC) | BsmI (GCATTC) | TAATACG ACTCACT ATA | - | - |
| **tRNA$^{Ile}$ ΔNCCA** | XbaI (TCTAGA) | SphI (GCATGC) | BsmI (GCATTC) | TAATACG ACTCACT ATA | - | - |
| **tRNA$^{Trp}$** | XbaI (TCTAGA) | SphI (GCATGC) | BsmI (GCATTC) | TAATACG ACTCACT ATA | - | - |

**Supplementary Table VIII. Relevant cloning details.**

### Supplementary Method

#### Fitting of biolayer interferometry data

Three binding models were considered to describe the binding kinetics: a one-step binding model, a two-independent binding sites or parallel one-step binding model, and a sequential two-step binding model. In the first model the binding describes one T-box molecule binding to one tRNA molecule following first order kinetics with one association and one dissociation constant (**Supplementary Figure 1C**). The parallel one-step binding model assumes two independent binding sites in the T-box and a tRNA binding to each site and is described by two sets of association and dissociation constants (**Supplementary Figure 1C**). Finally, the two-step binding model, described previously in a different context (3), assumes two binding sites between the T-box and the tRNA, and sequential binding to one site (Tbox:tRNA$_1$) followed by binding to the second site (Tbox:tRNA$_2$) (**Supplementary Figure 1C**). In this model, unbinding also assumes a sequential model but the first step is in only one direction due to the dilution of the tRNA.

The BLI data were fit by the three models using either the Octet Data Analysis Software version 11.1 (FortéBio Inc. San Jose, CA) for the one-step and parallel one-step binding models, or custom written software for the two-step binding model. The one-step and parallel one-step models did not fit the data well (**Supplementary Figure 2A**), while the two-step binding model produced the best results and described the binding process well (**Supplementary Figure 2A**). For that reason, all data were subsequently analyzed using the two-step binding model as described below.

In the two-step binding model, the dissociation kinetics were analyzed according to the following scheme:

$$Tbox : tRNA_2 \underset{k_{-2}}{\overset{k_2}{\rightleftharpoons}} Tbox : tRNA_1 \underset{k_{-1}}{\rightharpoonup} Tbox + tRNA$$

Which can be described by the following differential equations:

$$\frac{\partial[Tbox\colon tRNA_1](t)}{\partial t} = k_{-2}\,[Tbox\colon tRNA_2] - k_{-1}\,[Tbox\colon tRNA_1] - k_2\,[Tbox\colon tRNA_1]$$

$$\frac{\partial Tbox\colon tRNA_2](t)}{\partial t} = k_2\,[Tbox\colon tRNA_1] - k_{-2}\,[Tbox\colon tRNA_2]$$

The fitted parameters from the dissociation kinetics were then used to fit the following association kinetics according to the following scheme:

$$Tbox + tRNA \underset{k_{-1}}{\overset{k_1}{\rightleftharpoons}} Tbox : tRNA_1 \underset{k_{-2}}{\overset{k_2}{\rightleftharpoons}} Tbox : tRNA_2$$

Which can be described by the following differential equations:

$$\frac{\partial[Tbox\colon tRNA_1](t)}{\partial t} = k_1 \cdot [tRNA] \cdot [Tbox] - [Tbox\colon tRNA_1](k_1 \cdot [tRNA] + k_{-1} + k_2) - [Tbox\colon tRNA_2](k_1 \cdot [tRNA] - k_{-2})$$

$$\frac{\partial[Tbox\colon tRNA_2](t)}{\partial t} = k_2\,[Tbox\colon tRNA_1] - k_{-2}\,[Tbox\colon tRNA_2]$$

Association data were fitted by constraining $k_1$ to be smaller than the diffusion value (3) and the effective concentration of T-box in the biosensors to values previously reported for this type of biosensors (4).

The $K_D$ was calculated according to the formula (3):

$$K_D = \frac{k_{-1}}{k_1\left(1 + \frac{k_2}{k_{-2}}\right)}$$

Python scripts, along with the preprocessed data used to analyze these experiments, are available in the Source Data file.
